# Hijacking host microPEP: pathogens modulate the microRNA-microPEP loop to promote infection

**DOI:** 10.1101/2025.08.01.668066

**Authors:** Emmanuel Clostres, Christophe Penno, Abdelhak El-Amrani, Virginie Daburon, Kévin Gazengel, Cécile Monard, Stéphanie Daval

## Abstract

The partners of an ecological association tend to copy the biological system of their hosts. We hypothesized that microorganisms, particularly pathogens, have acquired the ability to express short peptides (pathoPEPs) homologous to host microPEPs (miPEPs) thus modulating the expression of the corresponding host microRNA (miRNA) and the function of miRNA-targeted genes. The pathosystem involving interactions between *Brassica napus* and its pathogen *Plasmodiophora brassicae* was studied. Using *in silico* analysis and ribosomal profiling, we identified three putative pathoPEPs produced by *P. brassicae* and their targeted plant miRNA genes. A link between the level of infection of *B. napus* by *P. brassicae* and the expression of pathoPEPs and their targeted miRNA genes was found, with the expression of the latter two being inversely related. Finally, we identified differential expression and translation of genes predicted to be targets of pathoPEP-regulated miRNAs. These genes, involved in auxin pathway, immune defense, root architecture or carbohydrate metabolism, are thought to enable *P. brassicae*, through its pathoPEPs, to hijack plant’s metabolic pathways (hormonal pathways, sugar synthesis, root morphology), thereby facilitating its invasion. Using computational *in silico* approaches, the involvement of miPEPs from plant pathogens as a host post-transcriptional regulatory pathway is described herein for the first time.

## INTRODUCTION

Micro ribonucleic acids (miRNAs) are small, non-coding RNAs typically 20-24 nt (see Krol et al., 2010 and Aguilar et al., 2019b for biogenesis). They play essential roles in regulating gene expression at the post-transcriptional level (Friedman et al., 2009; Tarver et al., 2012). They achieve this through two main mechanisms: firstly, by decreasing gene expression via translational repression, which lowers protein levels, and secondly, by cleaving target mRNAs (Borges and Martienssen, 2015). A specific gene expression pattern may then be involved in responses to biotic or abiotic stress.

Plant miRNAs have been reported to be involved in growth, development, and defense processes against pathogens (Kamthan et al., 2015; Luo et al., 2024). Numerous studies have demonstrated the functions of miRNAs in plant-microbe interactions and responses to infection, enabling plants to utilize their miRNAs to regulate their gene expression for pathogen defense. The roles of functionally validated miRNAs in mediating responses to various pathogens across different plant species at the molecular level are well described (Yang et al., 2021). Some plant miRNAs are known to regulate defense mechanisms against pathogenic infections, thereby boosting plant resilience against phytopathogens (Xie et al., 2017; Islam et al., 2018; Cui et al., 2020). For instance, the levels of the miR169 family members are correlated with susceptibility to fungal infections in plants such as *Musa acuminate* (Song et al., 2018). Additionally, evidence indicates that certain miRNAs contribute to pathogens resistance in the rice - *Magnaporthe oryzae* system: miR160 and miR398 act as positive regulators of immunity, while miR167 and miR319 serve as negative regulators (Asadi and Millar, 2024). Although the role of plant miRNAs in plant-microbe interactions has been extensively investigated across various pathosystems, there are limited reports detailing the specific functions of miRNAs in the interactions between Brassicas and *Plasmodiophora brassicae*, which causes clubroot disease, one of the biggest threats to rapeseed worldwide. Research has found evidence of miRNA-mediated regulation of genes controlling various developmental and stress-related traits in oilseed Brassicas (Rani et al., 2023). In *B. napus*, infection by *P. brassicae* alters the plant miRNA pool and target transcripts, regardless of the line’s susceptibility or resistance, or the disease development stage (Verma et al., 2014; Li et al., 2021). In *B. rapa*, miR1885s negatively modulate resistance genes in response to *P. brassicae* infection affecting disease tolerance (Paul et al., 2021). Similarly, plant miR167 negatively regulates *Arabidopsis thaliana* genes involved in immunity against *P. brassicae* (Liao et al., 2023).

Beyond to cellular, tissue, and inter-organ regulation within the same organism, miRNAs have emerged as crucial players in interkingdom communication, serving as functional messengers that bridge different species (Leitão et al., 2020; Jiang et al., 2023; Middleton et al., 2024; Chiquito-Contreras et al., 2024). These miRNAs are typically encapsulated within membranous vesicles known as exosomes (Choi et al., 2017) and our recent lab research has demonstrated their presence in exosomes released from the roots of *A. thaliana* and wheat (Penno et al., *in prep.*). The target organisms incorporate these encapsulated miRNAs which can regulate specific gene expressions, influencing related biological functions. Plants can produce specific miRNAs in response to infections, employing them as a direct defense against pathogens, similar to RNA interference. For instance, cotton plants release miR166 and miR159 to combat *Verticillium dahliae* infection by targeting the mRNA of fungal virulence factors. When these plant miRNAs bind to complementary sequences in the fungal mRNA, they trigger an RNA interference - like mechanism that degrades the targeted transcripts, thus diminishing *V. dahliae*’s virulence (Zhang et al., 2016). A comparable defensive strategy has been observed in wheat infected by *Fusarium graminearum*, where plant miR1023 silences an alpha/beta hydrolase gene linked to the fungus’s virulence (Jiao and Peng, 2018). Such discoveries suggest a widespread defensive mechanism in plants, where miRNAs target specific genes in the transcriptome of fungal pathogens to silence them (Asadi and Millar, 2024). Conversely, miRNAs may act as effectors from pathogens, modifying host immunity and infection dynamics by silencing genes in the host (Weiberg et al., 2014; Weiberg and Jin, 2015; Wang and Wang, 2016; Aguilar et al., 2019a; Arslan and Ozkilinc, 2024).

Interestingly, a recent discovery has revealed a subtle retro-control mechanism of miRNAs (Figure 1). Initially identified in the plant kingdom (Lauressergues et al., 2015) and later observed in human and animal systems (Ormancey et al., 2023), this mechanism involves micropeptides (miPEPs) composed of a few amino acids. These miPEPs are encoded by the primary transcript of mature miRNAs. Most investigations have reported that miPEPs up-regulate the expression of their corresponding primary transcripts (Lauressergues et al., 2022), but recent research (Ormancey et al., 2023) indicated that miPEPs can also down-regulate the expression of their corresponding miRNA primary transcript. Although the underlying mechanisms have not yet been identified, miPEPs may have different ways of affecting the expression of their corresponding miRNA primary transcript. Interestingly, recent data indicated that the mammalian gut microbiome expresses many small PEPs (Petruschke et al., 2021). Even though their functions have not been functionally analyzed, the authors suggested that they could be involved in microbiome-host communication.

**Figure 1:**
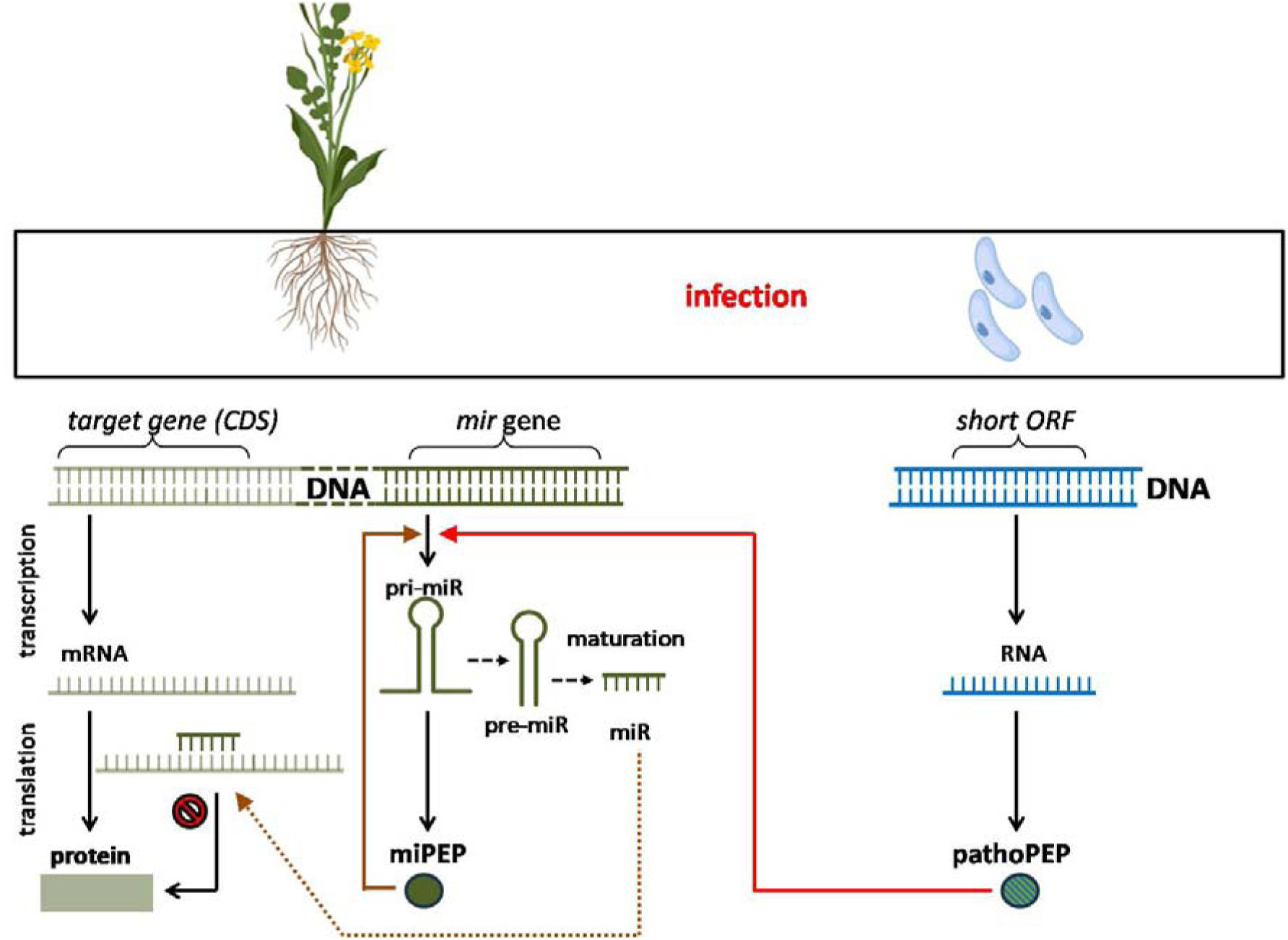
Retro-control mechanism of the plant microRNA-microPEP loop by the pathogen.

We hypothesize that the pathogen produces small PEPs homologues to hijack the host’s miRNA loop, thereby controlling host development and fitness. To enhance clarity, we will denote the peptides originated from the plant as miPEPs and those from the pathogen as pathoPEPs to highlight their microbial origin. To explore this potential communication pathway through miRNA and miPEP regulation in plant-pathogen interactions, we conducted a comprehensive *in silico* study alongside ribosome profiling (Ribo-seq) for the first time, using *B. napus* (rapeseed) *- P. brassicae* pathosystem as a model. Our goal was to predict and describe (i) plant-derived miPEPs and translated pathoPEPs, and (ii) targeted host miRNA along with their target genes.

## RESULTS

### Description of the pathosystem dataset and corresponding gradient of infection

Our computational prediction of pathoPEPs was based on RNA-seq data from the *B. napus* - *P. brassicae* pathosystem, gathered at different stages of clubroot infection (Daval et al., 2020) (Figure 2). The disease indexes were recorded as 19.79 ± 4.54%, 59.52 ± 3.48%, and 89.58 ± 1.04% for low, moderate, and severe infection levels, respectively. This equated to a *P. brassicae* DNA proportion in the roots of 0.04 ± 0.02, 0.29 ± 0.08, and 0.95 ± 0.06, respectively.

**Figure 2.**
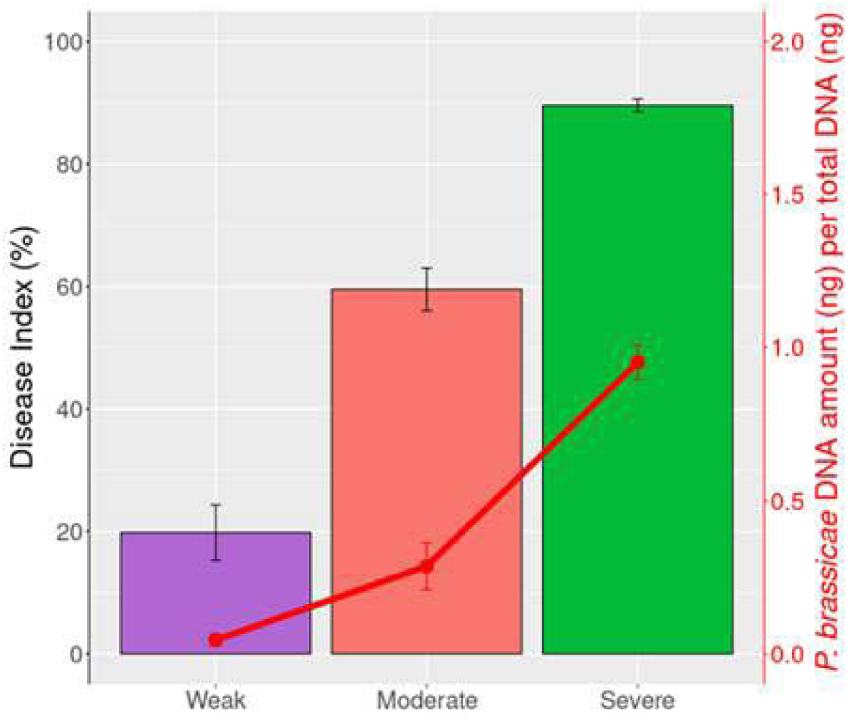
Gradient of clubroot development intensity (adapted from Daval et al. (2020)). Weak: *B. napus* Yudal 28 days, post inoculation with *P. brassicae*; Moderate: *B. napus* Tenor, 28 days post inoculation; Severe: *B. napus* Tenor, 36 days post inoculation. Clubroot symptoms were assessed using the disease index (histograms) and quantification of *P. brassicae* DNA through qPCR (curve), expressed as a ratio of the 18S DNA amount to total DNA. The data are the means of three biological replicates (12 plants per replicate), with error bars representing the standard errors of the means.

### Prediction of putative pri-miRNAs and their encoded miPEPs from *B. napus*

The prediction of pri-miRNAs from the genome of *B. napus* was conducted in two steps (Figure 3A). In the first step, as the pri-miRNAs were not annotated in existing miRNA databases, we searched the pre-miRNAs of *B. napus* in RNAcentral, identifying 2,253 pre-miRNAs. Additionally, we extracted 17,406 lncRNAs from the *B. napus* genome, along with gene features from available databases. After aligning the pre-miRNAs and lncRNAs datasets, we identified 51 putative pri-miRNAs derived from the lncRNA dataset, corresponding to 44 pre-miRNAs in RNAcentral. The presence of more pri-miRNAs than pre-miRNAs can be attributed to the alternative splicing of some lncRNA genes and the occurrence of certain pre-miRNAs across multiple lncRNA genes. In the second step, we hypothesized that the expression of miRNAs in *B. napus* is regulated by pathoPEPs produced by *P. brassicae*, prompting us to identify the differentially expressed pri-miRNAs of *B. napus* in the presence or absence of *P. brassicae*. Utilizing the gene expression dataset from Daval et al. (2020), we found 11, 38, and 32 pri-miRNAs that showed differential expression along the infection gradient, categorized as weak, moderate, and severe infection levels, respectively (Figure 3B). Since we anticipated that the expression of these pri-miRNAs would be regulated by pathoPEPs from *P. brassicae*, their sequences should contain the corresponding miPEP encoding sequences. Consequently, we predicted all potential miPEP sequences within the differentially expressed pri-miRNAs at each infection level, yielding to 271, 611, and 480 potential miPEPs, ranging from 6 to 100 amino acids, encoded by *B. napus* pri-miRNA at weak, moderate, and severe infection levels, respectively (Figure 3A).

**Figure 3.**
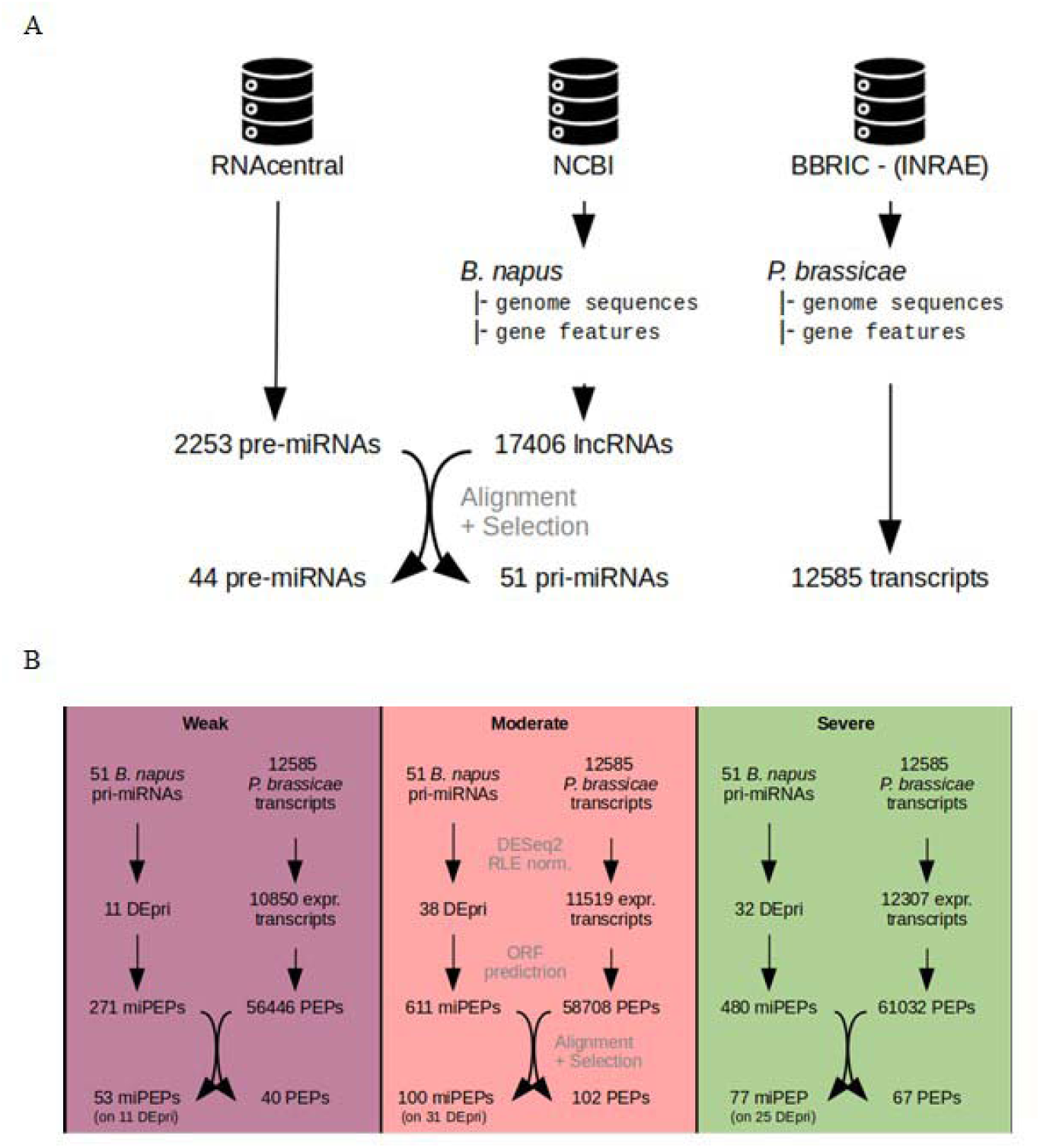
Number of *B. napus* pri-miRNAs and *P. brassicae* transcript sequences predicted and extracted from different databases (A) and number of pri-miRNAs, transcripts, miPEP and pathoPEP sequences obtained at each analysis step, categorized under weak, moderate and severe infection conditions (B).

### Prediction of pathoPEPs from *P. brassicae* and their targeted pri-miRNAs in *B. napus*

A total of 12,585 transcripts were retrieved from the *P. brassicae* genome. Of these, 10,850, 11,519, and 12,307 were expressed at weak, moderate, and severe infection levels according to the RNA-seq dataset of Daval et al. (2020), respectively (Figure 3B). From the expressed transcripts, potential pathoPEP sequences ranging from 6 to 100 amino acids were predicted, resulting in 56,446, 58,708, and 61,032 possible pathoPEPs from *P. brassicae* at weak, moderate, and severe infection levels, respectively. For each level of infection, these potential pathoPEPs were aligned with miPEPs encoded by pri-miRNA sequences from *B. napus* to identify pathoPEP-miPEP pairs. After this filtering step, the size distribution of pathoPEPs and miPEPs was reduced to a maximum of 18 amino acids. Final counts indicated 11, 31, and 25 pri-miRNAs of *B. napus* possibly regulated by 40, 102, and 67 predicted pathoPEPs from *P. brassicae* (totaling 106 distinct pathoPEPs), correlating to 7, 24, and 18 miRNA genes of *B. napus* (totaling 24 distinct miRNA genes) at weak, moderate, and severe infection levels, respectively (Figure 3B).

### Changes in the expression level of genes coding *B. napus* miRNAs and *P. brassicae* pathoPEPs depending on the infection level

From the RNA-seq dataset of Daval et al. (2020), we extracted the expression profiles of 24 *B. napus* miRNA genes which may be regulated by the predicted pathoPEPs of *P. brassicae*. Our differential expression analysis showed that these miRNA genes were downregulated in the presence of *P. brassicae* compared to the controls without the pathogen, regardless of the infection level. Meanwhile, the distribution pattern of all differentially expressed *B. napus* genes was similar for both upregulation and downregulation (Figure S1). Interestingly, the downregulation of the identified miRNA genes was more pronounced with increasing infection levels (Figure 4A). Conversely, the expression analysis of the genes encoding the predicted pathoPEPs of *P. brassicae*, which are believed to regulate the expression of *B. napus* miRNA gene, indicated that all of these genes were expressed, with their expression levels increasing in line with the infection severity (Figure 4B).

**Figure 4.**
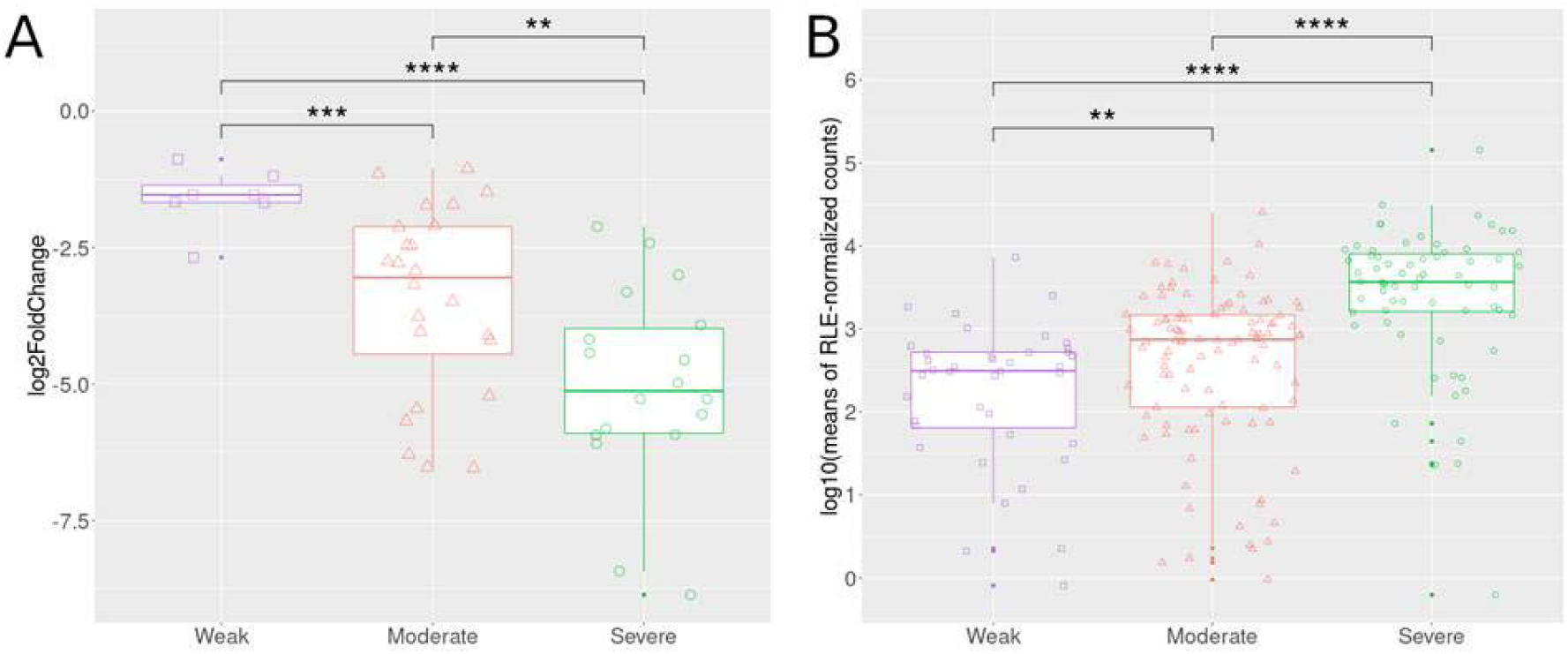
Variation in miPEP and pathoPEP gene expression across weak, moderate and severe infection levels. A: Differential expression of miRNA genes in *B. napus* targeted by the pathoPEPs from *P. brassicae* in uninfected *versus* infected plants. B: Expression of *P. brassicae* genes that code for pathoPEPs targeting miRNA genes in *B. napus*. Reads counts were normalized using Relative Log Expression (RLE). Asterisks (two to four) indicate significant differences in the means of log2FoldChange (A) or normalized counts (B) between the two conditions compared, with p-value of less than or equal to 1e^-2^, 1e^-3^ or 1e^-4^, respectively.

### Interactions between the target miRNA genes of *B. napus* and the predicted pathoPEPs from *P. brassicae* - constancy along the gradient of infection

Among the 24 differentially expressed genes (DEGs) in *B. napus* that correspond to the identified miRNA, four were found across all three infection levels. Likewise, from the 106 expressed genes in *P. brassicae* that encode the predicted pathoPEPs, 28 were shared among the three infection levels (Table S1). In detail, we observed that (i) at the weak level, 40 pathoPEPs targeted 7 genes specifying for 11 pri-miRNAs, yielding 2 mature miRNAs; (ii) at the moderate level, 102 pathoPEPs targeted 24 genes specifying for 31 pri-miRNAs, resulting in 10 mature miRNAs; (iii) at the severe level, 67 pathoPEPs targeted 18 genes specifying for 25 pri-miRNAs, yielding to 8 mature miRNAs. Thus, several pathoPEPs can target the same pri-miRNA, while different pri-miRNAs can yield the same mature miRNA.

### Ribosome footprint of the candidate pathoPEPs

To investigate the reality of a miRNA-pathoPEP regulation loop, a ribosomal profiling approach was conducted using samples from experimental growth of the oilseed rape genotypes Tenor and Yudal, both inoculated with the eH isolate of *P. brassicae*.

We validated our experimental setup by quantifying *P. brassicae* in the two genotypes of *B. napus* using qPCR. As anticipated, the infection level in Tenor was significantly greater than that in Yudal (P<0.05; Figure S2). We analyzed the reads generated from both Ribo-seq (Figure S3) and RNA-seq, with the former mapping 6 to 10% (6-11 million reads) and the latter 58 to 62% (14-15 million reads) to the concatenated transcriptome of *B. napus* and *P. brassicae*, respectively. The low proportion of mapped reads from Ribo-seq resulted from a significant number of reads aligning to rRNA, despite the performed rRNA depletion (52-58 % of the raw reads) and a notable number of duplicate reads (26-30 % of the raw reads).

All 106 expressed genes from the *in silico* analysis of pathoPEPs were predicted on coding transcripts. The high ribosomal coverage of coding sequences hindered the clear identification of translation signals for candidate pathoPEPs in different frames from the coding DNA sequence (CDS). We focused on 27 pathoPEPs predicted in untranslated regions (UTRs), including 5 in the 5’UTR and 22 in the 3’UTR (Table S2). Twelve pathoPEPs in UTRs showed ribosome footprints in at least one replicate, with three selected (Pldbra_eH_r1s016g07830_1, Pldbra_eH_r1s019g08936_2, Pldbra_eH_r1s020g09116_2) for further analysis due to their localization outside annotated CDSs (Figure S4). Among these, Pldbra_eH_r1s016g07830_1 exhibited strong reproducibility, showing a ribosome footprint in all three biological replicates of Tenor genotype. In contrast, ribosomal coverage for Pldbra_eH_r1s020g09116_2 and Pldbra_eH_r1s019g08936_2 was observed in only one replicate. Pldbra_eH_r1s020g09116_2 was the only putative pathoPEP consistently detected across all three infection levels. Pldbra_eH_r1s019g08936_2 was detected at both moderate and severe infection levels, whereas Pldbra_eH_r1s016g07830_1 was exclusively observed at the moderate infection level.

### *B. napus* genes targeted by the pathoPEP-regulated miRNAs

We identified three distinct *B. napus* miRNAs specified in a pri-miRNA sequence that encodes a miPEP homologous to one of the pathoPEPs found by Ribo-seq. The miRNA (pathoPEP) pairs were miR156 (Pldbra_eH_r1s020g09116_2), miR158 (Pldbra_eH_r1s019g08936_2), and miR172 (Pldbra_eH_r1s016g07830_1). Among them, only miR156 was differentially expressed between the two plant genotypes, showing significantly higher expression in Yudal compared to Tenor (p = 0.0023, logFC = −3.2). To rule out the possible synthesis of plant miPEPs from these three miRNA genes, we investigated their possible translation. No ribosome footprints were detected, indicating that these miRNA genes did not produce plant miPEPs homologous to the putative pathoPEPs at the time of sampling.

We investigated the *B. napus* genes targeted by miR156, miR158, and miR172, analyzing both RNA-seq and Ribo-seq datasets to identify differentially expressed genes (DEGs) and differentially translated genes (DTGs) between the two genotypes. We identified 17, 5, and 10 DEGs targeted by miR156, miR158, and miR172, respectively. Of these, 10, 2, and 6 genes were significantly more expressed in Tenor than in Yudal (Figure S5A). At the translational level, DTGs were observed only among the genes targeted by miR156 (10 DTGs) and miR158 (5 DTGs), with 6 and 2 of these genes, respectively, being over-translated in Tenor compared to Yudal (Figure S5B). In total, 13 genes were both differentially expressed and translated between both genotypes: 8 genes targeted by miR156 and 5 by miR158.

We then examined the functions of all DEGs and DTGs in *B. napus*. Functional annotation was not available for 12 DEGs and 6 DTGs. Notably, among the DEGs and DTGs targeted by miR156, which were expressed more in Tenor than in Yudal, we identified four functions that were also translated more in Tenor. These functions included a glucuronoxylan 4-O-methyltransferase 3 (GXM3; FDR = 0.030, logFC = 0.741 and FDR = 0.048, logFC = 0.869 in RNA-seq and Ribo-seq datasets, respectively), an auxin response factor 6 (ARF6; FDR = 0.001, logFC = 1.234 and FDR = 0.025, logFC = 1.228 in RNA-seq and Ribo-seq datasets, respectively), a putative nuclear RNA export factor SDE5 isoform X3 (FDR = 0.001, logFC = 1.234 and FDR = 0.005, logFC = 1.175 in RNA-seq and Ribo-seq datasets, respectively), and a squamosa promoter-binding-like (SPL) protein 2 (FDR = 0.001, logFC = 2.717 and FDR = 0.027, logFC = 2.252 in RNA-seq and Ribo-seq datasets, respectively). In contrast, protein 6 of this SPL transcription factor was translated more in Yudal than in Tenor (FDR = 0.050 and logFC = −1.392). Among the five genes targeted by miR158 that were differentially expressed and translated, only two had functional annotation. In Tenor, the potential hexosyltransferase gene, MUCI70, showed significantly higher expression and translation than in Yudal (FDR = 0.018, logFC = 1.329; FDR = 0.033, logFC = 1.611 in RNA-seq and Ribo-seq datasets, respectively). In contrast, a transcription factor, MYB20, which is encoded by two different genes (LOC106394697 and LOC106411208), was the only annotated function that was significantly more expressed and translated in Yudal compared to Tenor (FDR = 0.001/0.001, logFC = −3.481/−2.434; FDR = 0.046/0.039, logFC = −1.755/−1.814 in RNA-seq and Ribo-seq datasets, respectively).

## DISCUSSION

### Can the pathogen produce pathoPEPs homologous to the plant’s miPEPs?

Small RNAs (sRNAs), particularly miRNAs, play crucial roles in the regulation of plant basal defense and immunity (Navarro et al., 2006; Li et al., 2014). For example, infection by *P. brassicae* alters the miRNA pool and target transcripts in the host plant (Li et al., 2021). In response, many pathogens have developed the capability to generate their own sRNA-like molecules that mimic plant sRNA, facilitating their infection. A case in point is the wheat pathogen *Puccinia striiformis*, which produces miRNA-like RNAs that act as effectors to diminish host immunity (Wang et al., 2017). Given that the expression of plant miRNA is regulated by pri-miRNA-encoded peptides, pathogens may take advantage of this mechanism by producing mimicry effectors, referred to as pathoPEPs, which specifically regulate the synthesis of related host miRNA influencing the immune response both directly and indirectly. Small peptides have been recognized as significant modulators of plant immunity, enhancing or fine-tuning immune signaling, thereby raising the possibility of pathogen exploitation of this system (Segonzac and Monaghan, 2019). Furthermore, administering specific miPEPs to *A. thaliana* has demonstrated the ability to upregulate the expression of their corresponding pri-miRNA, yielding to either enhanced or diminished expression of genes involved in plant’s response to various pathogens, which corresponds to reduced infection or increased susceptibility, respectively (Ormancey et al., 2024). These findings prompt us to hypothesize that plant pathogens might produce pathoPEPs to manipulate the host’s miRNA synthesis, ultimately enhancing the silencing or induction of gene expression to facilitate their infection process.

To test our hypothesis, we combined data from Daval et al. (2020) on the *B. napus* – *P. brassicae* pathosystem with computational predictions and ribosome profiling. As previously highlighted (Makarewich and Olson, 2017), both methodologies have been fine-tuned to identify small ORFs encoding miPEPs. From this published paper, we utilized transcriptomic data (RNA-seq) alongside the infection levels in two genotypes of *P. brassicae*-infected and healthy *B. napus*. Although the infection gradient emerged from RNA-seq data of two genotypes, Yudal (exhibiting weak infection) and Tenor (showing moderate to severe infection levels), we anticipated to identify shared pathoPEPs and miRNA genes across all infection levels. This expectation relies on utilizing a reference genome from another *B. napus* genotype to predict potential pri-miRNAs and their associated miPEPs, which are believed to exist independently of the genotype.

From these data, we predicted pathoPEP genes in the *P. brassicae* genome and identified their target miRNA genes in *B. napus*. We found that the expression of pathoPEP genes increased with greater infection severity, while the expression of their predicted target host miRNA genes decreased. In contrast to the positive feedback regulation typically associated with plant miPEPs (Lauressergues et al., 2022), our findings suggest that pathoPEP may repress the expression of corresponding host miRNA genes. A similar down-regulation of miRNA gene expression caused by miPEP has been observed in mice, where miPEP31 acts as a transcriptional repressor of miRNA-31 (Zhou et al., 2022). Therefore, the function of miPEP does not appear to be universally conserved as previously observed by Prel et al. (2021).

### Are the predicted pathoPEPs actually translated?

The regulation of *B. napus* genes through pathoPEP required effective translation. To advance beyond the *in silico* prediction of pathoPEPs, a ribosome profiling approach was performed in both Tenor and Yudal *B. napus* genotypes to determine which of the 106 pathoPEPs predicted *in silico* are actively translated. Our analysis focused on three translated pathoPEPs located outside an annotated coding sequence and their potential role in regulating *B. napus* gene expression. This Ribo-seq approach provides insights into actively translated mRNA by capturing transcripts shielded within ribosome complexes. When paired with RNA-seq, this approach enables a thorough analysis of both transcriptional and translational changes that could differ between sensitive (Tenor) and resistant (Yudal) plant genotypes.

Key considerations must be emphasized regarding the limitations of Ribo-seq in identifying translated sequences. Firstly, higher transcription levels of certain coding genes provide more mRNA templates for translation, resulting in a greater yield of representative ribosome-protected fragments (RPFs) compared to coding sequences with lower transcription levels. Secondly, larger coding sequences can accommodate more ribosomes than shorter ones. Since RPFs typically range from 26 to 34 nt in size, larger coding sequences may result in RPF populations displaying overlapping sequence identities based on the ribosomes’ positions on the sequences. Conversely, as we included PEP ranging from 10 to 100 amino acids, this means that a 10 amino-acid peptide is encoded by a 30 nt sequence, allowing for at most one ribosome-protected fragment. As a result, translating short PEPs from short miPEP and pathoPEP coding sequences is likely to generate a less diverse range of RPFs. Third, since *P. brassicae* requires the infection of the plant root, the preparation of Ribo-seq was conducted from infected plants, which contained cells from both the plant and the pathogen. Therefore, the prevalence of background sequences plays a significant role in identifying low-abundant sequences that encode pathoPEPs while preparing Ribo-seq samples. Thus, caution should be exercised when interpreting Ribo-seq results focuses on identifying pathoPEPs, particularly given the potential bias associated with their detection due to their possibly low expression and limited sequence length for ribosome protection alongside the ratio of infected to uninfected host cells present in the sample mixture used for Ribo-seq preparation.

### Which host miRNAs are targeted and regulated by translated pathoPEPs?

The three identified pathoPEPs target miR156, miR158 and miR172 in *B. napus*. Interestingly, the translation of the putative pathoPEP Pldbra_eH_r1s020g09116_2 was associated with the repression of its target gene miR156 in the susceptible Tenor genotype, which correlated with heightened infection severity. In *B. napus* infected with *P. brassicae*, miR156 was shown to be regulated depending on the stage of infection and to be involved in hormone homeostasis (Verma et al., 2014). Generally, mechanisms of clubroot susceptibility or resistance have been shown to be controlled by miRNAs and their target plants during *P. brassicae* infection (Wei et al., 2023). Collectively, this provides further evidence of a potential hijacking of the plant miRNA pathway by the pathogen.

### Which host functions are potentially controlled by the pathogen through pathoPEP-regulated miRNAs?

The goal of pathoPEPs is to regulate host gene expression and protein synthesis by modulating the expression levels of host miRNAs. Thus, we aimed to highlight the biological functions of *B. napus* regulated by miRNAs that are directly targeted by pathoPEPs. By employing Ribo-seq and RNA-seq, we identified differential expression and translation of genes predicted to be targets of miRNAs (miR156, miR158 and miR172) regulated by pathoPEP across two *B. napus* genotypes, reflecting changes linked to infection level. We expected that *P. brassicae* would more effectively enhance its infection strategies in the susceptible Tenor genotype compared to the resistant Yudal genotype. For miR172, we found that targeted genes were either differentially expressed or translated, but not both. In contrast, we identified target genes for miR156 and miR158 that were expressed and translated at significantly higher levels in Tenor compared to Yudal. Interestingly, the target genes for mir156 include a glucuronoxylan 4-O-methyltransferase 3 (GXM3), a SQUAMOSA promoter binding protein-like (SPL), a putative mRNA export factor (SDE5), and an auxin response factor (ARF6), while those for mir158 include a glycosyltransferase (MUCI70) (Figure 5).

**Figure 5.**
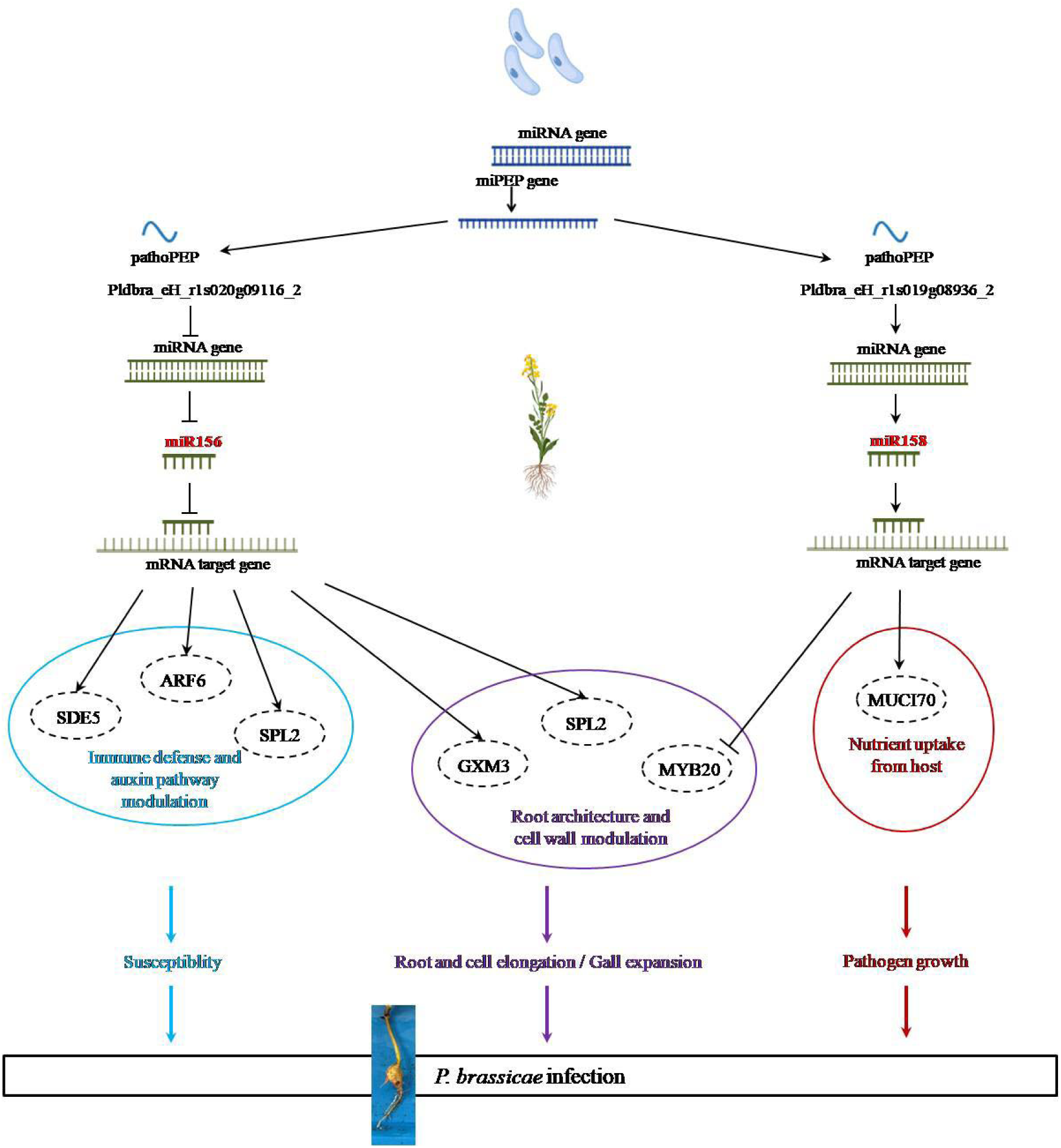
Proposed model of *P. brassicae* infection by hijacking the host plant microRNA-microPEP loop through its pathoPEPs.

Among the genes targeted by miR156, we identified GXM3 encoding for a glucuronoxylan 4-O-methyltransferase 3, an enzyme involved in the xylan biosynthesis (Liang et al., 2019), secondary cell walls synthesis, and plant growth (Lee et al., 2012). Previously, GXM3 was identified as a miR146 target in *Brassica campestris* (Liang et al., 2019). In our experiment, the downregulation of mir156 in Tenor led to an upregulation and enhanced translation of the GXM3 gene. The dynamic alterations of cell walls in Brassicas during clubroot disease are well described, with the disease marked by the formation of root galls causing changes and reorganization of the cell wall (Badstöber et al., 2020). The overexpression of the GXM3 gene could contribute to the formation of hypertrophied roots upon *P. brassicae* infection, providing novel insights into how *P. brassicae* uses these modifications to benefit its growth and development.

Another gene targeted by mir156 correspond to SPL encoding for a SQUAMOSA promoter binding protein-like 2, which belongs to a family of transcription factors regulated by miR156 (Rhoades et al., 2002; Xu et al., 2016; Sang et al., 2023; Mehtab-Singh et al., 2024). SPL proteins, which are specific to plants, have overlapping but distinct functions in promoting adult characteristics, including leaf morphogenesis, floral induction, and reduced adventitious root development in *A. thaliana* (Wu et al., 2009). Although there are several SPL genes, SPL2 is particularly important for vegetative development (Xu et al., 2016). It is worth noticing that the overexpression of miR156 decreases the levels of SPL2 mRNAs (Kasschau et al., 2003; Vazquez et al., 2004; Schwab et al., 2005). This aligns with our study, where overexpressed miR156 led to significantly lower SPL2 expression in Yudal compared to the susceptible Tenor genotype. Interestingly, recent studies have shown that the miR156-SPL pathway can also be involved in plant immunity. For instance, in *Salvia miltiorrhiza*, SPL2 overexpression modified root architectures by affecting auxin biosynthesis and signaling (Wang et al., 2022). In *A. thaliana*, disrupting miR156 function compromised jasmonic acid and salicylic acid signaling, and disease resistance to *Botrytis cinereac* (Sun et al., 2022) and to *Pseudomonas syringae* (Yin et al., 2019). By decreasing miR156 expression and so increasing SPL2 gene expression, pathoPEP Pldbra_eH_r1s020g09116_2 may hinder root growth while altering the plant’s hormonal response, facilitating infection.

Another gene that miR156 targets is SDE5 which encodes a putative mRNA export factor. SDE5 plays a crucial role in silencing transgenes and generating trans-acting siRNAs. Its expression is associated with plant defense responses. Notably, transgenic *A. thaliana* plants that overexpress SDE5 display a weakened defense response (Uddin et al., 2017). Furthermore, mutation in SDE5 also results in increased susceptibility to cucumber mosaic virus in *A. thaliana* (Hernandez-Pinzon et al., 2007). In our study, the reduced miR156 expression in Tenor compared to Yudal led to the upregulation of both transcription and translation of SDE5, which could potentially reduce the immune response against *P. brassicae* infection.

The fourth gene of interest targeted by miR156 is ARF6, which encodes an auxin response factor. Auxin plays a crucial role in regulating various aspects of plant growth and development, and is also significant in responding to pathogens and disease progression. It regulates gene expression through ARF family transcription factors that activate or repress target genes based on specific DNA binding domain. The repression of the auxin signaling pathway is known to contribute to plant immune defenses, downregulation of this pathway enhances resistance against pathogens (Wang et al., 2007; Llorente et al., 2008; Fu and Wang, 2011). This is particularly notable with biotrophs pathogen like *P. brassicae*, as up-regulation of auxin signaling is linked to greater host susceptibility to diseases (Kunkel and Johnson, 2021), leading to heightened symptoms such as gall development (Vañó et al., 2023) and promoting host cells expansion (Ludwig-Müller and Schuller, 2008), and modulation of pathways associated with auxin biosynthesis, homeostasis, or receptors after clubroot infection (Robin et al., 2020).

ARF6, a member of the ARF transcription factors, was shown to be overexpressed and overtranslated in the Tenor genotype susceptible to infection (Li et al., 2016), which can help *P. brassicae* development in two major ways. First, ARF6 regulates processes like root development and osmotic stress responses (Gutierrez et al., 2009), which the pathogen hijacks to facilitate its entry. Secondly, the ARF6 gene is the target of miR156 which could control both the auxin pathway and immune response. While Gutierrez et al. (2009) identified ARF6 as a target of miR167, they overlooked miR156, despite miRNAs’ known role in auxin-mediated plant responses to pathogens (Navarro et al., 2006). A link between the auxin pathway and miR167 has been found, particularly with its target gene ARF8. For example, in *A. thaliana* infected with *Pseudomonas syringae*, miR167 is expressed differently in response to the pathogen, and its overexpression enhance resistance by suppressing auxin responses, partially relying on salicylic acid signaling (Caruana et al., 2020). Similarly, during *A. thaliana’ s* infection by *P. brassicae*, miR167 was inhibited, resulting in high expression level of one of its target genes, ARF8 (Liao et al., 2023). Just as miR167 may affect the plant’s response to clubroot by regulating ARF8, miR156 could influence the auxin pathway, the homeostasis of plant hormones, and the response to clubroot disease by regulating ARF6. In our study, the elevated expression and translation levels of ARF6 in the susceptible Tenor genotype may reflect a pathoPEP-mediated mechanism that *P. brassicae* uses to weaken immune resistance, likely by enhancing auxin signaling to inhibit defense pathways and boost host susceptibility.

Finally, one of the genes of interest predicted to be targeted by miR158 is MUCI70, which encodes for a glycosyltransferase that belongs to the carbohydrate-active enzymes (CAZymes) family (Lombard et al., 2014) (database http://www.cazy.org/, v12), which was overexpressed and overtranslated in Tenor, the susceptible genotype. During *P. brassicae* infection, altered plant primary and secondary metabolism is known, including pathways involved in carbohydrates (Li et al., 2020). The published genomes for *P. brassicae* exhibit typical biotrophic features of obligate biotrophic pathogens. This includes a reduced number of CAZymes that are involved in the synthesis, metabolism, and transport of carbohydrates, as well as incomplete carbohydrate biosynthetic pathways, suggesting a dependency on nutrient intake from the host (Schwelm et al., 2015; Rolfe et al., 2016; Bi et al., 2016; Daval et al., 2019). It is described that *P. brassicae* recruits plant SWEET sucrose transporters within developing galls to assimilate host plant sugars (Walerowski et al., 2018). Additionally, through the overexpression of MUCI70 regulated by miR156, itself regulated by pathoPEP, *P. brassicae* could reprogram carbohydrate production for its own benefit, facilitating local distribution of sugars to itself, and producing a sink of plant metabolites that it can assimilate. This active metabolic pathway hijacking via pathoPEP allows *P. brassicae* to take up nutrients from the host cells and facilitates its invading progression.

The second gene of interest also targeted by miR158 is MYB20. This was the only gene that was significantly more expressed and translated in the partially resistant Yudal genotype compared to Tenor. MYB20 is a transcription activator of genes involved in lignin and phenylalanine biosynthesis during secondary wall formation in *A. thaliana* (Geng et al.), and may aid to immunity against necrotrophic pathogens (Lai and Mengiste, 2013). In potatoes infected with *Pectobacterium carotovorum* ssp. *brasiliense*, MYB20 was also down-regulated in the susceptible cultivar (Kwenda et al., 2016). The activation of lignin-related pathways is generally linked with clubroot resistance, with resistant varieties showing lignin accumulation around the penetration site (Tu et al., 2024). Reduced MYB20 levels in the susceptible genotype may impair plant growth and structural reinforcement of the cell wall during *P. brassicae* infection, suggesting that the *P. brassicae* pathoPEPs may disrupt lignin activation and hinder a robust plant resistance mechanism.

## CONCLUSION

This pioneering work within a pathosystem provides novel insight into how the pathogen might regulate the expression of its host’s miRNAs by hijacking its miPEPs regulation system for miRNA gene expression and producing mimicked miPEPs, the pathoPEPs. Taken together, these findings suggest several pathways potentially regulated by pathoPEPs through miRNA-mediated modulation of gene expression and translation. These pathways encompass not just immune responses but also essential developmental processes tied to plant growth and structural integrity. In the *B. napus* – *P. brassicae* pathosystem, this dual influence may serve as a strategic approach for the pathogen to undermine host defenses while fostering an environment conducive to its own growth. Understanding this molecular basis of plant-microbe-pathogen interactions is important for advancing agricultural sustainability.

## METHODS

### Computational *in silico* prediction of PEPs from plant (miPEPs) and pathogen (pathoPEPs), and their plant miRNA targets

#### Plant-microorganism models and data sources

The pathosystem *B. napus* (rapeseed) and its bioaggressor *P. brassicae* eH (protista) serves as a valuable model for investigating the potential involvement of miPEPs and pathoPEPs in this interaction, bolstered by available genomics and experimental transcriptomics data for both partners (Daval et al., 2019; Daval et al., 2020). The overall workflow for *in silico* prediction of PEPs from plant (miPEPs) and pathogen (pathoPEPs) is illustrated in Figure 6. The genome (FASTA) and gene features (GFF3) for the *B. napus* cultivar “ZS11” (GCF_000686985.2) and *P. brassicae* eH (POCA00000000.1) were downloaded from the NCBI and BBRIC (INRAE, Toulouse, France) databases, respectively. Transcriptomic data for *B. napus* and *P. brassicae* were obtained from Daval et al. (2020) and are available in the European Nucleotide Archive database. These data correspond to the simultaneous sequencing of both organisms (dual RNA-seq) extracted from the plant roots of two *B. napus* genotypes, Tenor and Yudal, which displayed different susceptibilities to clubroot infection by *P. brassicae*, with Tenor being more susceptible than Yudal. To investigate gene expression in *B. napus* and *P. brassicae*, their genomes, annotation files, and transcriptome reads from the roots of *B. napus* (both uninfected and infected by *P. brassicae*) were imported into Galaxy (https://usegalaxy.org/).

**Figure 6.**
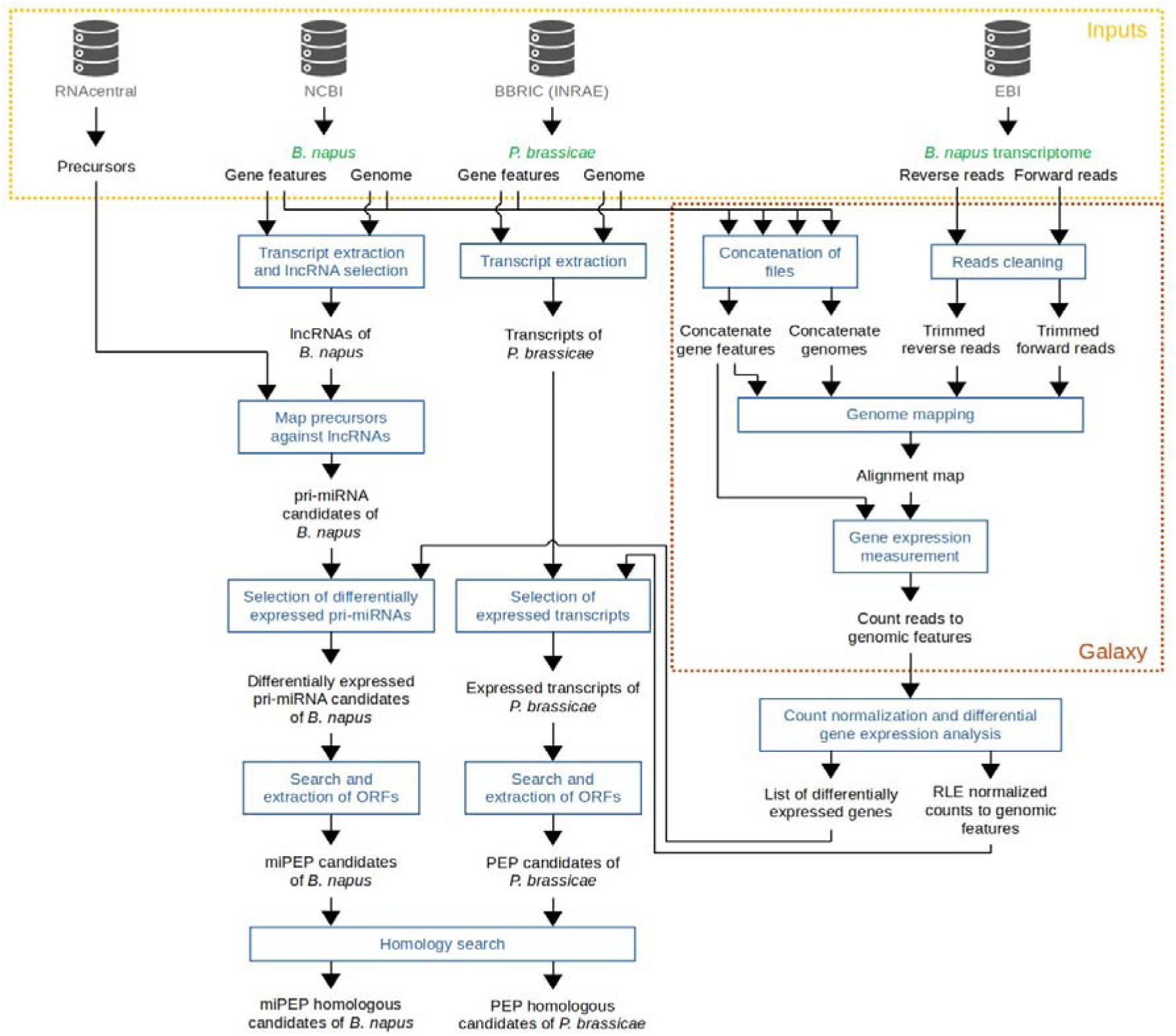
Homemade workflow for identifying plant miPEPs and pathogen pathoPEPs. In gray are databases; green, biological materials; blue, analyses performed; black, the input/output files of each analysis.

Since the pri-miRNAs of *B. napus* are not annotated in the existing miRNA databases, we initially focused on the long non-coding RNA (lncRNA) containing miRNA precursor (pre-miRNA) sequences obtained from the RNAcentral database (Sweeney et al., 2021). This database is distinctive as it consolidates miRNAs annotated in other databases, providing more potential miRNA precursors than alternatives like miRBase. According to our hypotheses, the anticipated features of the *B. napus*’s pri-miRNAs include: (i) the presence of a precursor miRNA sequence listed in RNAcentral, (ii) differential expression levels in infected plants compared to uninfected ones, depending on the infection level, and (iii) inclusion of PEP open reading frames (ORFs) that are homologous to the *in silico* predicted pathoPEPs in *P. brassicae*.

#### Transcriptomic data from B. napus and P. brassicae pathosystem

Transcriptomic reads were recovered from the roots of infected plants under three experimental conditions representing gradual levels of infection: weak, moderate, and severe (Figure 2). These data correspond to three biological replicates (each composed of 12 pooled plants) of Yudal or Tenor genotypes sampled 28 days after inoculation (weak and moderate infection levels, respectively) and of Tenor genotype sampled 36 days after inoculation (severe infection level). Similarly, the transcriptomic data from corresponding uninfected plants were fetched as a control.

#### Identification of putative pri-miRNAs of B. napus from pre-miRNAs and transcriptomic data (Figure 6)

All transcripts of *B. napus* and *P. brassicae* were extracted from the genome sequences (FASTA) and gene features (GFF3) using AGAT v0.9.1 (Dainat et al., 2022). For *P. brassicae*, ribosomal RNAs were excluded. For *B. napus*, only transcripts annotated as “lncRNAs” were retained (Bartel, 2004). To predict putative pri-miRNAs, a semi-global alignment was conducted using glsearch36 v36.3.8i (Pearson, 2016) between the pre-miRNAs and the lncRNAs. The alignment was global on pre-miRNAs and local on lncRNAs. Putative pri-miRNAs were identified as lncRNAs that present a perfect alignment (no substitutions, no gaps) with the precursors.

#### Selection of differentially expressed pri-miRNAs of B. napus and expressed transcripts of P. brassicae

The genomes and annotation files for both *B. napus* and *P. brassicae* were combined using Concatenate datasets v0.1.1 tool from Galaxy wrapper. Reads were cleaned with Trim Galore v0.6.7 (https://github.com/FelixKrueger/TrimGalore accessed July 18th, 2022) to remove sequences with a phred score ≤ 25 and a length < 50 nt. The cleaned reads were subsequently aligned with the concatenated files using STAR v2.7.8a (Dobin et al., 2013). Mapped reads were then assigned to genomic features and counted using featureCounts v1.6.3 (Liao et al., 2014).

After filtering out read counts equal to 0 across the three biological replicates, the remaining read counts were normalized using the Relative Log Expression (RLE) method in the DESeq2 v1.36.0 package (Love et al., 2014) in R (version 4.2.1; R Core Team, 2022). A differential expression analysis was performed on the normalized counts, retaining only those differentially expressed genes (DEGs) with a False Discovery Rate (FDR) ≤ 0.01, specifically between the *P. brassicae* infected and uninfected conditions.

The list of DEGs in *B. napus* was used to identify differentially expressed pri-miRNAs (DEpri). log2FoldChange (LFC) differing from zero indicated miRNA genes that were differentially expressed between the conditions of *P. brassicae* infection and no infection. Pri-miRNAs with LFC > 0 were categorized as upregulated miRNA genes, while those with LFC < 0 were deemed downregulated miRNA genes in infected plants compared to uninfected ones.

Considering that *P. brassicae* cannot survive independently of its plant host, conducting differential gene expression analysis of the pathogen between free-living and plant-associated states was not possible. Consequently, we selected transcripts with a normalized RLE count > 0, which correspond to transcripts expressing detectable RNA-seq at the three levels of infection.

#### Search for predicted miPEPs and pathoPEPs and analysis of their homologies

All potential candidate, miPEPs from *B. napus* and pathoPEPs from *P. brassicae*, were retrieved using getorf (from EMBOSS v6.6.0.0; Rice et al., 2000) to extract coding sequences (CDSs) that encode peptides ranging from 6 to 100 amino acids. These CDSs were exclusively predicted based on the forward strand of the differentially expressed predicted pri-miRNAs (DEpri) in *B. napus* (being the source of miPEP) and the expressed transcripts of *P. brassicae*.

To identify the pathoPEPs that might regulate the expression of host plant pri-miRNAs, a global alignment was conducted between candidate pathoPEPs and plant miPEPs using ggsearch36 v36.3.8i. Homologous pathoPEP-miPEP pairs were defined as those exhibiting at least 50% identity, a maximum of 2 gap openings, and an alignment e-value of 0.01 or lower.

### Experimental investigation of the putative pathoPEP-miRNA loop using ribosomal profiling

#### Experimental setup, plant material, pathogen inoculation and disease assessment

The protocols for obtaining and characterizing the samples used in Ribo-seq are described in File S1.

#### Sample preparation, RNA sequencing, ribosome profiling and bioinformatic analyses

EIRNA Bio (https://eirnabio.com) generated RNA-seq and ribosomal profiling (Ribo-seq) libraries from frozen root samples of infected Tenor (n=3) and Yudal (n=3) genotypes. The experimental procedure involved gently lysing the cell membrane to release its contents, which were then subsampled for Ribo-seq and RNA-seq. Ribo-seq enables the identification of translated RNA sequences (Ingolia et al., 2009). 90% of the lysate volume was used for Ribo-seq to isolate polysomes associated with RNA transcripts, followed by RNase treatment to yield monosomes containing ribosome-protected fragments (RPF) of 26-34 nt. Next, RNA isolation was carried out, along with size selection of the 26-34 nt fragments. To remove rRNA contaminants from the RNA population, ten custom hybridization probes were used, and Ribo-seq analysis was performed on all 2 x 3 samples. The remaining 10% of the lysate was used for total RNA extraction and the preparation of counterpart RNA-seq libraries to identify the transcriptome from which the RPF fragments are derived. Both Ribo-seq and RNA-seq libraries were sequenced on Illumina’s Nova-seq 6000 platform, targeting approximately 100 million reads per sample for Ribo-seq and between 20 to 30 million for RNA-seq. The NGS reads are deposited in the European Nucleotide Archive (ENA) database system (EBI) under accession numbers ERS25060339 to ERS25060350 (project PRJEB90664; https://www.ebi.ac.uk/ena/browser/view/PRJEB90664).

Reads quality was assessed using FastQC (version 0. 11. 9) (https://www.bioinformatics.babraham.ac.uk/projects/fastqc/), and adapters were trimmed using Cutadapt v 5. 0 (Martin, 2011) with settings tailored for small reads analysis in Ribo-seq. Reads were then mapped to the *B. napus* and *P. brassicae* genomes using the STAR aligner (Dobin et al., 2013).

To enhance the likelihood of locating translated pathoPEPs, the Ribo-seq data of the Tenor genotype, which had the highest infection level, was analyzed. This involved searching for ribosome footprints exhibiting a read count greater than 0 for each putative pathoPEP on the transcripts encoding them in *P. brassicae*. Only pathoPEPs displaying a translation signal, and not co-localized with an annotated CDS (regardless of reading frame) were retained for further analysis.

#### Identification of differentially expressed and translated plant pri-miRNA genes regulated by the pathoPEPs and their targeted genes

The analysis of DEGs for miR genes potentially regulated by pathoPEPs, along with the analysis of differential expression and translation of genes targeted by these miR were performed between the Yudal and Tenor genotypes, which exhibit significantly different infection levels (Figure S2). The askoR R package was utilized for these analyses (https://github.com/askomics/askoR). By filtering out low-count genes, only those with at least 0.5 counts per million reads in at least three samples were retained. DEGs were identified through a generalized linear model utilizing likelihood ratio tests (glmLRT in edgeR v.3.40.2), and the Trimmed Mean of M-values (TMM) normalization method was applied. Genes were deemed differentially expressed if they had an FDR-adjusted p-value of less than 0.05, according to the Benjamini-Hochberg correction, without applying any logFC threshold filtering. Functional annotation of each targeted gene was performed using Blast2go version 1.5.1.

### Statistical analyses

Statistical analyses and figure generation were conducted using R (version 4.2.1) with the “stats” and “ggplot2” packages (Wickham, 2016). A p-value threshold of 0.01 was applied. The Shapiro-Wilk normality test (Royston, 1982) was utilized to assess normality before performing pairwise comparisons of means across the three infection levels. If the data met the normality assumption, we performed a t-test or a Welch t-test when the variances of the compared conditions were similar or different (as determined by the F test), respectively. For the data that did not meet the normality assumption, we used the Mann-Whitney test. These tests were executed either (i) on LFCs of *B. napus* DEpri encoding putative homologous miPEPs, or (ii) on normalized mean counts (using the RLE method) of expressed *P. brassicae* transcripts that encode putative homologous pathoPEPs. Disease data were analysed using a likelihood ratio test using a cumulative link model (CLMM; “clmm” function, “ordinal” package). LSMeans and pairwise comparisons of LSMeans were calculated with the Tukey test (α = 5%), using the ‘cld’ function of the ‘lsmeans’ package.

P-values from the tests were adjusted through the Benjamini-Hochberg (BH) method. One, two, three, or four asterisks “*” were assigned when the adjusted p-value was less than or equal to 5e^-2^, 1e^-2^, 1e^-3^, or 1e^-4^, respectively. If not significant, “NS” was indicated.

## ACKNOWLEDGMENTS

The authors thank Jean-Philippe Combier (Laboratoire de Recherche en Sciences Végétales, CNRS/UT3/INPT, 31320 Auzeville-Tolosane, France) for interactive communication, as well as Bruno Marquer (IGEPP, INRAE), Fabrice Legeai (IGEPP, INRAE), Quentin Le Galludec, and Anne-Solène Airiau for technical assistance. They also acknowledge the Biological Resources Center BrACySol (INRAE Rennes, France) for providing Brassica seeds and the molecular ecology platform (PEM, ECOBIO) for technical support.

## AUTHOR CONTRIBUTIONS

AE-A designed the research, supervised and revised the manuscript. EC, CP, AE-A, CM and SD designed the experiments. Bioinformatics analyses were performed by EC and KG. Ribo-seq analysis was performed by EC and CP. Molecular experiments were performed by VD. EC, CM, CP and SD wrote the manuscript. All authors read and approved the final manuscript.

---

**Figure S1.**
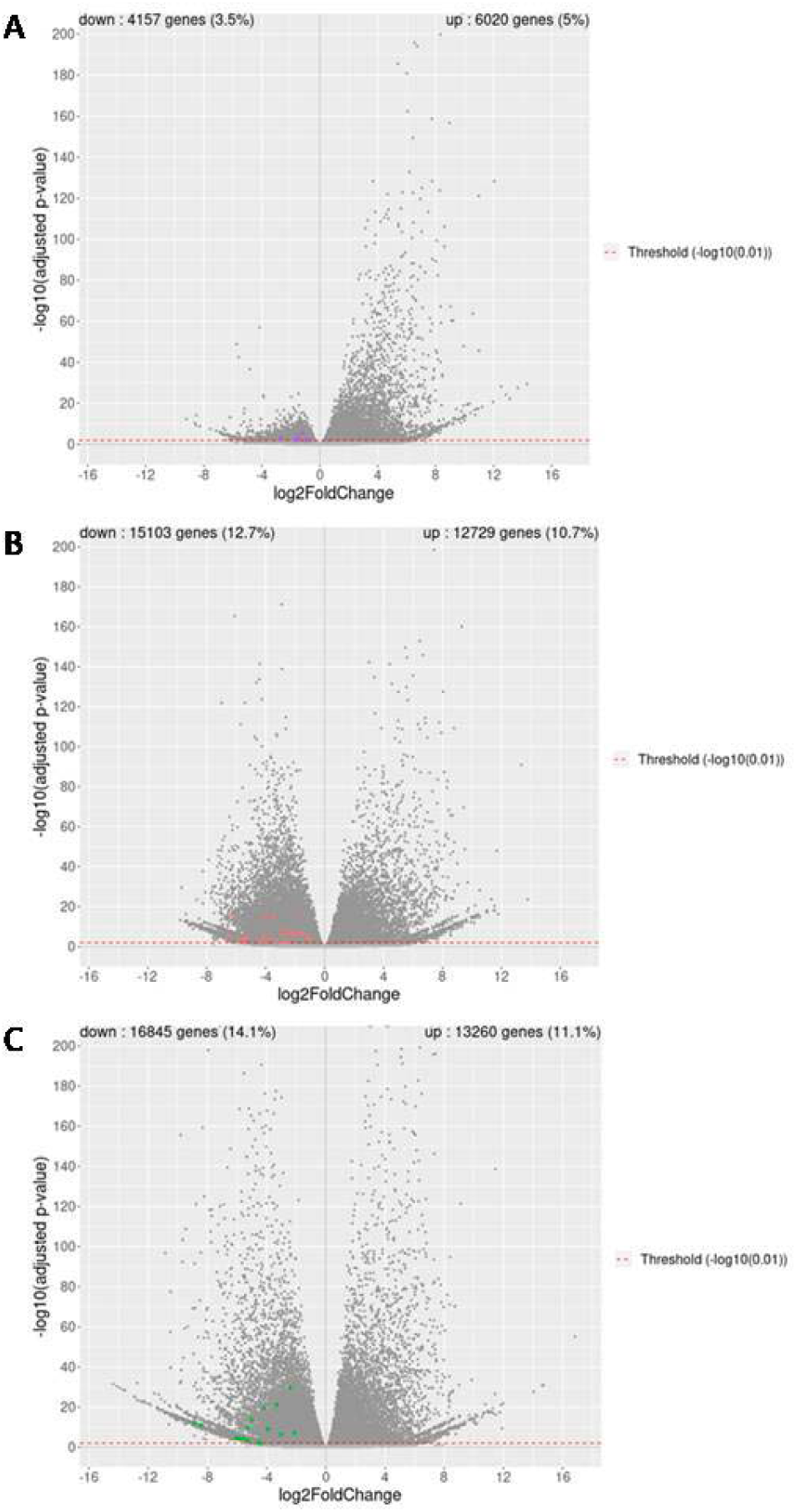
Volcano plot of differential gene expression of *B. napus* uninfected compared to weak (A), moderate (B), or severe (C) *P. brassicae* infection levels. Each point represents a gene, indicating its significance (p-value) relative to the log2FoldChange. These two parameters are calculated by DESeq2 using three biological replicates. A difference in expression is deemed significant for a p-value of 0.01 (red broken horizontal line). The 7, 24 and 18 miRNA genes of *B. napus* that are targeted by the pathoPEPs predicted from *P. brassicae* transcripts are shown in purple (A), light red (B) and green (C).

**Figure S2.**
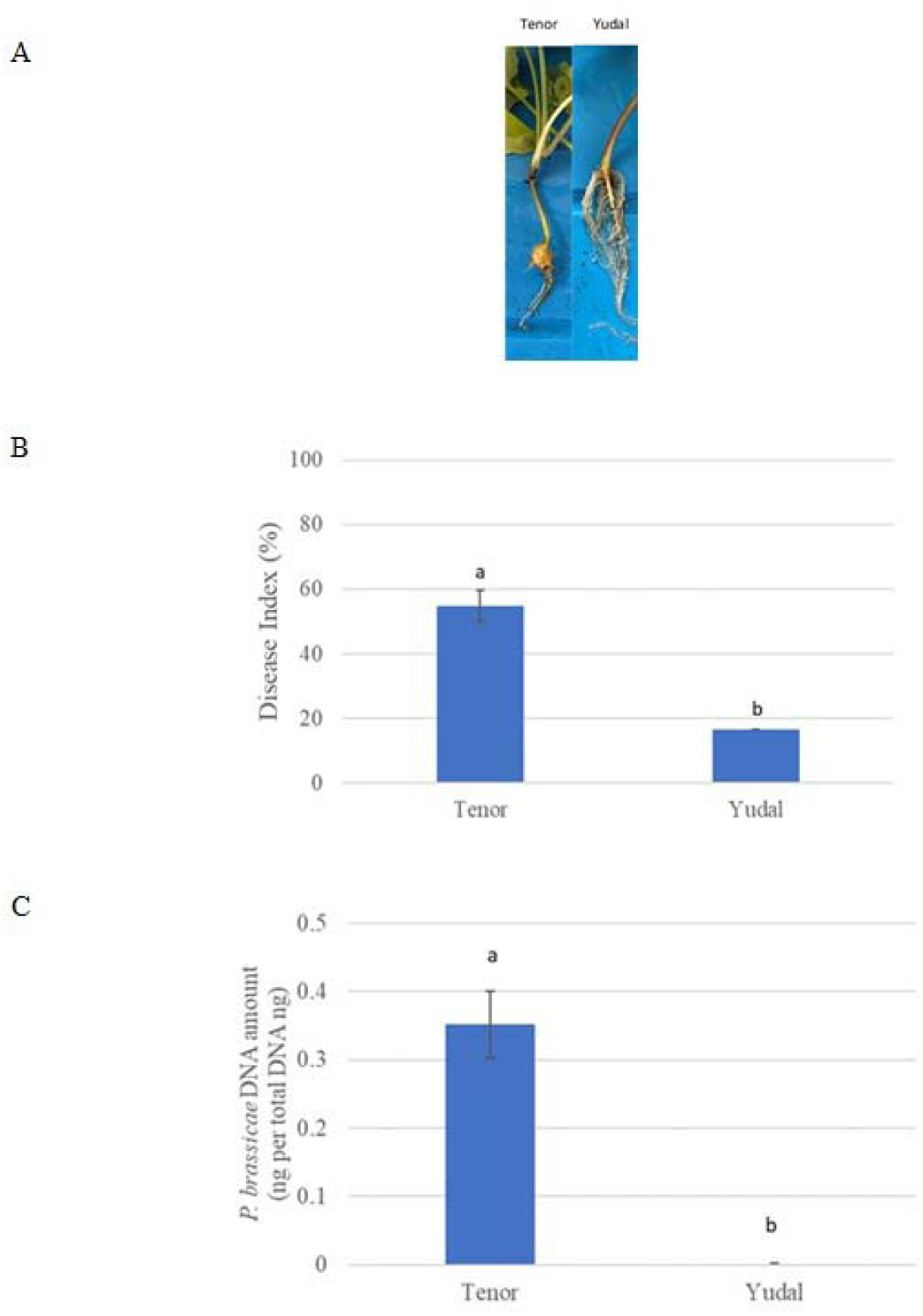
Clubroot development. Clubroot symptoms **(A)** were assessed using the disease index **(B)** and the quantification of *P. brassicae* DNA by qPCR **(C)**, calculated as a ratio of the 18S DNA amount to total DNA. The data represent the means of three biological replicates (12 plants per replicate), with error bars indicating standard errors of the means. Different letters denote statistically significant differences between genotypes at *P* < 0.05.

**Figure S3.**
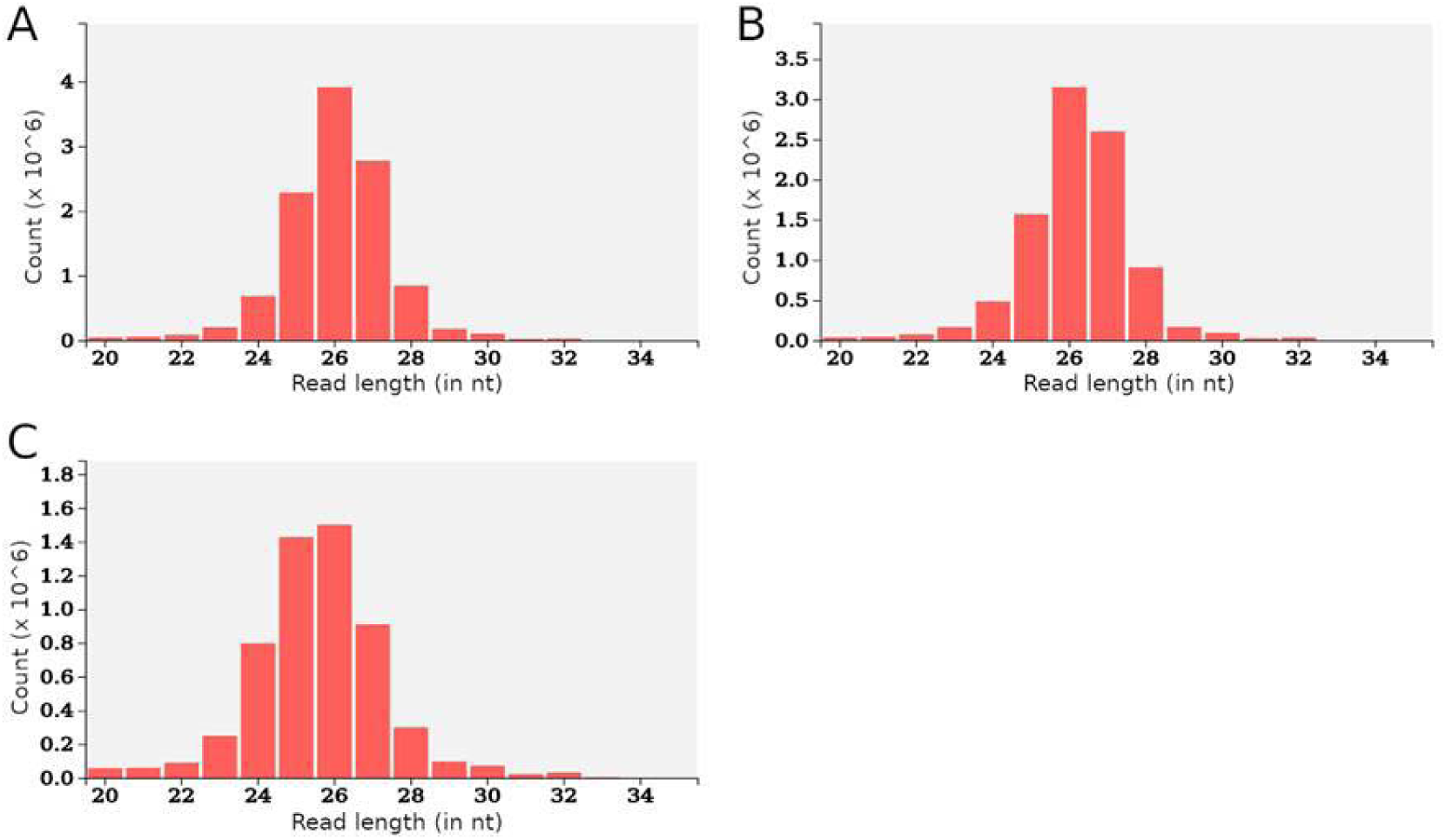
Distribution of the nucleotide length of ribosomal protected fragments derived from Ribo-seq reads for the three biological replicates of the infected Tenor genotype.

**Figure S4.**
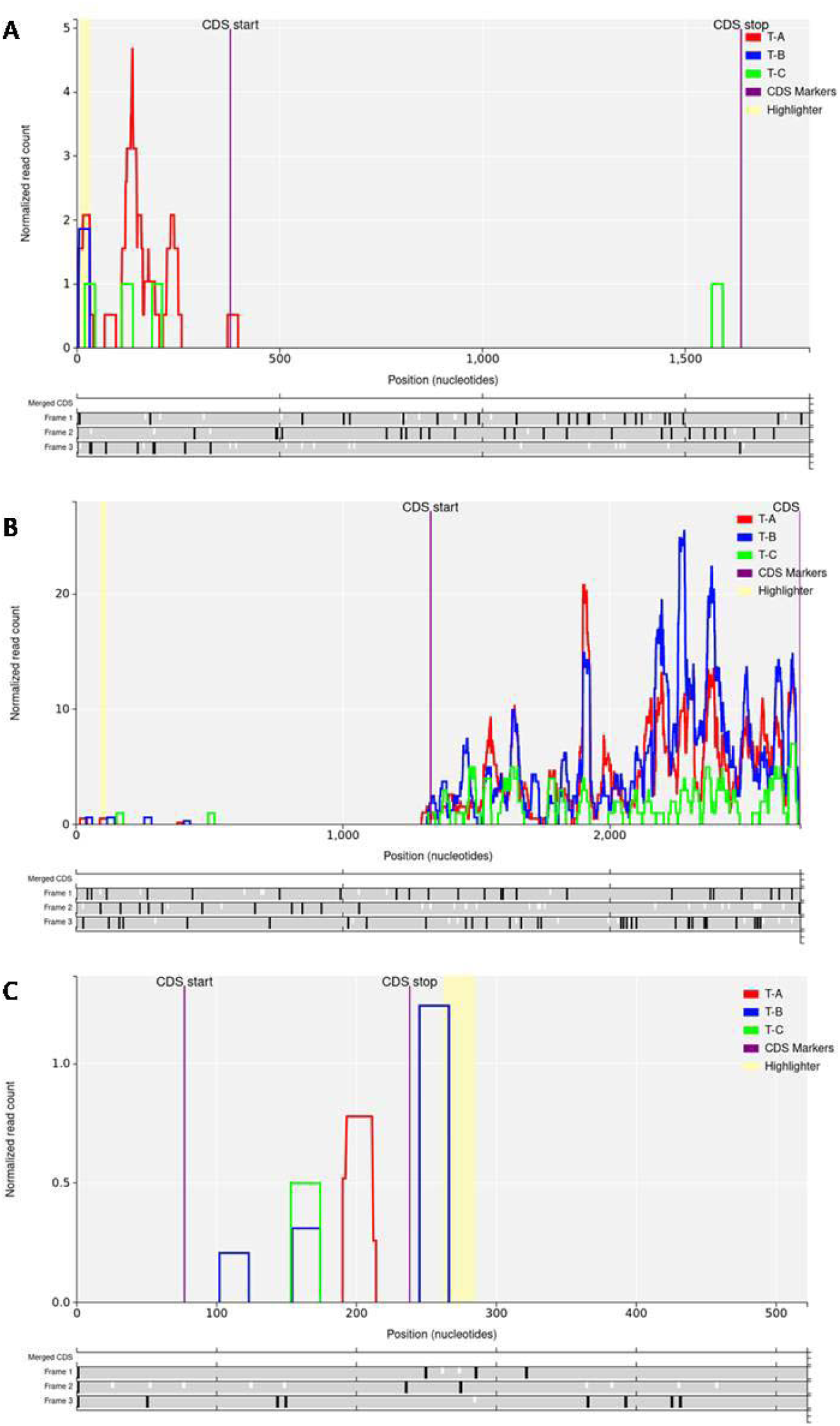
Plot of the ribosome footprint of pathoPEP candidate. Coverage of ribo-seq along the 3 frames of *P. brassicae* transcripts Pldbra_eH_r1s016g07830 **(A)**, Pldbra_eH_r1s020g09116 **(B)**, and Pldbra_eH_r1s019g08936 **(C).** T-A (red), T-B (blue), and T-C (green) are the biological replicates of *B. napus* Tenor infected with *P. brassicae*. Purple lines denote the start and stop codons of the CDS. The yellow highlighter indicates the position of the pathoPEP candidate in frame 3 of the transcript. Vertical black bars on the different frames mark canonical stop codons (UAA, UAG or UGA), while vertical white bars denote AUG start codons.

**Figure S5.**
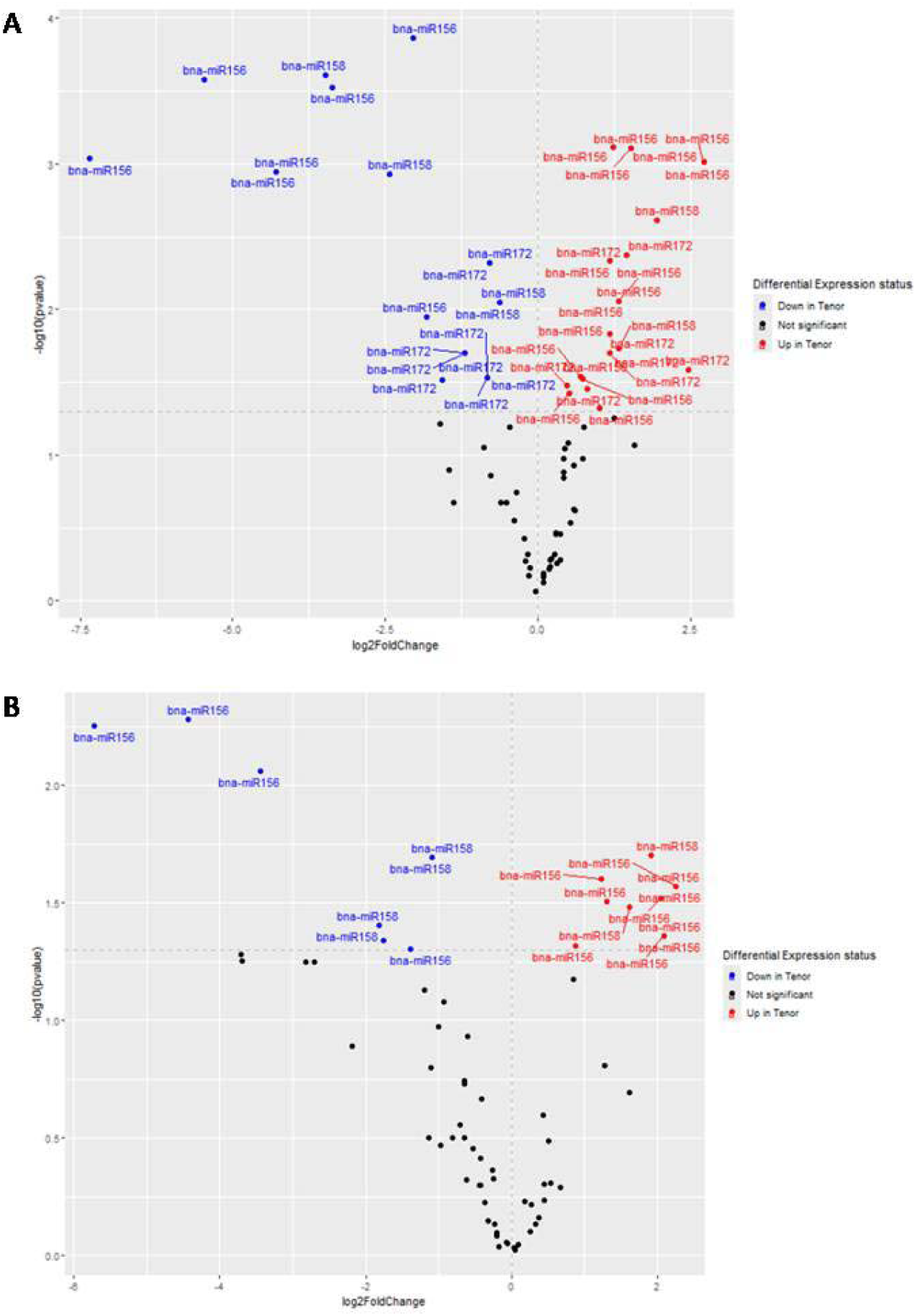
Comparative analysis of Yudal and Tenor genotypes. Volcano plots of **(A)** the differentially expressed genes DEGs, and (B) the differentially translated genes DTGs, targeted by the 3 miRNAs. The log2FC indicates the mean expression level for each gene. Each dot represents a single gene. Black dots indicate no significant DEGs or DTGs, blue dots represent down-regulated genes in Tenor compared to Yudal, and red dots indicate up-regulated genes in Tenor compared to Yudal.

**Table S1.** List of *P. brassicae* pathoPEPs targeting the miRNA genes of *B. napus* across the three infection levels.

**Table S2.** Position, region and ribosome coverage of the 27 pathoPEPs predicted in the untranslated regions (UTR). A “-” indicates no ribosomal coverage, while one to three “+” signs indicate ribosomal coverage observed in one, two, or three biological replicates, respectively.

## File S1

### File S1 Supplementary methods

#### Experimental design, plant material, pathogen inoculation, and disease assessment employed in Ribo-seq

The genotypes Tenor and Yudal of *B. napus*, along with the eH isolate of *P. brassicae* (pathotype P1) were utilized (Daval et al., 2019). *B. napus* seeds were sown in pots containing 400 g of soil and were incubated in a climatic chamber. Three replicates, each with 12 plants, were prepared for the two conditions (Yudal and Tenor inoculated with *P. brassicae*). The plants were inoculated with a resting spore suspension of the *P. brassicae* eH isolate. To prepare the inoculum, clubs grown on the universal susceptible host, Chinese cabbage (*B. rapa* ssp *pekinensis* cv. *Granaat*), were collected, blended with sterile water, and filtered through layers of cheesecloth. The resting spores were then filtered through 500, 100, and 55 µM sieves to remove plant cell debris. The spore concentration was determined using a Malassez cell and adjusted to 1 × 10[ spores ml^-1^. Plant inoculation involved treating ten-day-old seedlings by applying 1 ml of the spore suspension at a concentration of 1 × 10[ spores ml^-1^ to the base of each stem. The plants were maintained at 22°C during the day and 19°C at night, under a 16-hour photoperiod. They were periodically watered with a Hoagland nutritive solution to ensure nutrient supply and maintain a water retention capacity of 70– 100%. Roots from 12 plants were sampled 45 days after inoculation (dai).

For each replicate, eight plants were selected for disease assessment and pathogen quantification. First, clubroot symptoms were evaluated using a disease index based on localization and size of galls. Next, the roots were cut below the collar, at a depth of −1 to −6 cm and separated from the soil, then washed twice in sterile water for 10 s each time using vortexing. Afterward, the roots were cut into small pieces, frozen in liquid nitrogen, and stored at −80°C. After lyophilization, dry root biomass was measured, and the powder was retained for DNA extraction and pathogen quantification via qPCR (Daval et al., 2020).

Briefly, 1 µl of DNA extracted from the root samples was used for quantitative PCR on the LightCycler® 480 Real-Time PCR System (Roche) to quantify *P. brassicae*. This involved amplifying a 164 pb sequence of the 18S gene with primers: 5[-TTGGGTAATTTGCGCGCCTG-3[ (forward) and 5[-CAGCGGCAGGTCATTCAACA-3[ (reverse). Each qPCR reaction of 20 µl contained 10 µl of SYBR Green Master Mix (Roche), 0.08 µl of each primer (100 µM), and 1 µl of total DNA as the template. The PCR conditions consisted of an initial denaturation at 95°C for 5 min, followed by 45 cycles at 95°C for 10 s and 64°C for 40 s. Standard curves were generated using serial dilutions of *P. brassicae* DNA extracted from resting spores. Quantitative results were then expressed and normalized to reflect the mean *P. brassicae* DNA content in the total root-extracted DNA. For each replicate (n=3), roots from the four remaining plants were also sampled, washed in sterile water, cut into small pieces and frozen in liquid nitrogen prior to being stored at −80°C for further ribosomal profiling.

## REFERENCES

Aguilar, C., Mano, M., and Eulalio, A. (2019a). MicroRNAs at the host–bacteria interface: host defense or bacterial offense. Trends Microbiol. 27:206–218. 10.1016/j.tim.2018.10.011.

Aguilar, C., Mano, M., and Eulalio, A. (2019b). Multifaceted roles of microRNAs in host-bacterial pathogen interaction. Microbiol. Spectr. 7. 10.1128/microbiolspec.bai-0002-2019.

Arslan, K., and Ozkilinc, H. (2024). MicroRNA cross-talk between Monilinia fungal pathogens and peach host. Phytoparasitica 52:1–16. 10.1007/S12600-024-01131-Z.

Asadi, M., and Millar, A. A. (2024). Review: Plant microRNAs in pathogen defense: A panacea or a piece of the puzzle? Plant Sci. 341:111993. 10.1016/J.PLANTSCI.2024.111993.

Badstöber, J., Ciaghi, S., and Neuhauser, S. (2020). Dynamic cell wall modifications in brassicas during clubroot disease. bioRxiv Advance Access published March 3, 2020, doi:10.1101/2020.03.02.972901. 10.1101/2020.03.02.972901.

Bi, K., He, Z., Gao, Z., Zhao, Y., Fu, Y., Cheng, J., Xie, J., Jiang, D., and Chen, T. (2016). Integrated omics study of lipid droplets from Plasmodiophora brassicae. Sci. Reports 2016 61 6:1–12. 10.1038/SREP36965.

Borges, F., and Martienssen, R. A. (2015). The expanding world of small RNAs in plants. Nat. Rev. Mol. Cell Biol. 2015 1612 16:727–741. 10.1038/NRM4085.

Caruana, J. C., Dhar, N., and Raina, R. (2020). Overexpression of Arabidopsis microRNA167 induces salicylic acid-dependent defense against Pseudomonas syringae through the regulation of its targets ARF6 and ARF8 4:e00270. 10.1002/PLD3.270.

Chiquito-Contreras, C. J., Meza-Menchaca, T., Guzmán-López, O., Vásquez, E. C., and Ricaño-Rodríguez, J. (2024). Molecular insights into plant–microbe interactions: a comprehensive review of key mechanisms. Front. Biosci. - Elit. 16:9. 10.31083/j.fbe1601009.

Choi, J.-W. W., Um, J.-H. H., Cho, J.-H. H., and Lee, H.-J. J. (2017). Tiny RNAs and their voyage via extracellular vesicles: Secretion of bacterial small RNA and eukaryotic microRNA. Exp. Biol. Med. 242:1475–1481. 10.1177/1535370217723166.

Cui, C., Wang, J. J., Zhao, J. H., Fang, Y. Y., He, X. F., Guo, H. S., and Duan, C. G. (2020). A Brassica miRNA regulates plant growth and immunity through distinct modes of action. Mol. Plant 13:231–245. 10.1016/J.MOLP.2019.11.010.

Dainat, J., Hereñú, D., Davis, E., Crouch, K., LucileSol, Agostinho, N., pascal-git, and tayyrov (2022). AGAT Advance Access published 2022, doi:10.5281/ZENODO.6621429. 10.5281/ZENODO.6621429.

Daval, S., Belcour, A., Gazengel, K., Legrand, L., Gouzy, J., Cottret, L., Lebreton, L., Aigu, Y., Mougel, C., and Manzanares-Dauleux, M. J. (2019). Computational analysis of the Plasmodiophora brassicae genome: mitochondrial sequence description and metabolic pathway database design. Genomics 111:1629–1640. 10.1016/J.YGENO.2018.11.013.

Daval, S., Gazengel, K., Belcour, A., Linglin, J., Guillerm-Erckelboudt, A. Y., Sarniguet, A., Manzanares-Dauleux, M. J., Lebreton, L., and Mougel, C. (2020). Soil microbiota influences clubroot disease by modulating Plasmodiophora brassicae and Brassica napus transcriptomes. Microb. Biotechnol. 13:1648–1672. 10.1111/1751-7915.13634.

Dobin, A., Davis, C. A., Schlesinger, F., Drenkow, J., Zaleski, C., Jha, S., Batut, P., Chaisson, M., and Gingeras, T. R. (2013). STAR: ultrafast universal RNA-seq aligner. Bioinformatics 29:15–21. 10.1093/BIOINFORMATICS/BTS635.

Friedman, R. C., Farh, K. K. H., Burge, C. B., and Bartel, D. P. (2009). Most mammalian mRNAs are conserved targets of microRNAs. Genome Res. 19:92–105. 10.1101/GR.082701.108.

Fu, J., and Wang, S. (2011). Insights into auxin signaling in plant-pathogen interactions. Front. Plant Sci. 2:12287. 10.3389/fpls.2011.00074.

Geng, P., Zhang, S., Liu, J., Zhao, C., Wu, J., Cao, Y., Fu, C., Han, X., He, H., and Zhao, Q. MYB20, MYB42, MYB43, and MYB85 regulate phenylalanine and lignin biosynthesis during secondary cell wall formation. Plant Physiol. 182. 10.1104/PP.19.01070.

Gutierrez, L., Bussell, J. D., Pǎcurar, D. I., Schwambach, J., Pǎcurar, M., and Bellini, C. (2009). Phenotypic plasticity of adventitious rooting in Arabidopsis is controlled by complex regulation of AUXIN RESPONSE FACTOR transcripts and microRNA abundance. Plant Cell 21:3119–3132. 10.1105/TPC.108.064758.

Hernandez-Pinzon, I., Yelina, N. E., Schwach, F., Studholme, D. J., Baulcombe, D., and Dalmay, T. (2007). SDE5, the putative homologue of a human mRNA export factor, is required for transgene silencing and accumulation of trans-acting endogenous siRNA 50:140–148. 10.1111/J.1365-313X.2007.03043.X.

Ingolia, N. T., Ghaemmaghami, S., Newman, J. R. S., and Weissman, J. S. (2009). Genome-wide analysis in vivo of translation with nucleotide resolution using ribosome profiling. Science 324:218. 10.1126/SCIENCE.1168978.

Islam, W., Qasim, M., Noman, A., Adnan, M., Tayyab, M., Farooq, T. H., Wei, H., and Wang, L. (2018). Plant microRNAs: Front line players against invading pathogens. Microb. Pathog. 118:9–17. 10.1016/J.MICPATH.2018.03.008.

Jiang, C. H., Li, Z. J., Zheng, L. Y., Yu, Y. Y., and Niu, D. D. (2023). Small RNAs: Efficient and miraculous effectors that play key roles in plant–microbe interactions. Mol. Plant Pathol. 24:999–1013. 10.1111/MPP.13329.

Jiao, J., and Peng, D. (2018). Wheat microRNA1023 suppresses invasion of Fusarium graminearum via targeting and silencing FGSG_03101. J. Plant Interact. 13:514–521. 10.1080/17429145.2018.1528512.

Kamthan, A., Chaudhuri, A., Kamthan, M., and Datta, A. (2015). Small RNAs in plants: recent development and application for crop improvement. Front. Plant Sci. 06. 10.3389/fpls.2015.00208.

Kasschau, K. D., Xie, Z., Allen, E., Llave, C., Chapman, E. J., Krizan, K. A., and Carrington, J. C. (2003). P1/HC-Pro, a viral suppressor of RNA silencing, interferes with Arabidopsis development and miRNA function. Dev. Cell 4:205–217. 10.1016/S1534-5807(03)00025-X.

Krol, J., Loedige, I., and Filipowicz, W. (2010). The widespread regulation of microRNA biogenesis, function and decay. Nat. Rev. Genet. 2010 119 11:597–610. 10.1038/NRG2843.

Kunkel, B. N., and Johnson, J. M. B. (2021). Auxin plays multiple roles during plant-pathogen interactions. Cold Spring Harb. Perspect. Biol. 13. 10.1101/CSHPERSPECT.A040022.

Kwenda, S., Motlolometsi, T. V., Birch, P. R. J., and Moleleki, L. N. (2016). RNA-seq profiling reveals defense responses in a tolerant potato cultivar to stem infection by Pectobacterium carotovorum ssp. Brasiliense. Front. Plant Sci. 7:238216. 10.3389/FPLS.2016.01905.

Lai, Z., and Mengiste, T. (2013). Genetic and cellular mechanisms regulating plant responses to necrotrophic pathogens. Curr. Opin. Plant Biol. 16:505–512. 10.1016/J.PBI.2013.06.014.

Lauressergues, D., Couzigou, J.-M., Clemente, H. S., Martinez, Y., Dunand, C., Bécard, G., and Combier, J.-P. (2015). Primary transcripts of microRNAs encode regulatory peptides. Nature 520:90–93. 10.1038/nature14346.

Lauressergues, D., Ormancey, M., Guillotin, B., San Clemente, H., Camborde, L., Duboé, C., Tourneur, S., Charpentier, P., Barozet, A., Jauneau, A., et al. (2022). Characterization of plant microRNA-encoded peptides (miPEPs) reveals molecular mechanisms from the translation to activity and specificity. Cell Rep. 38:110339. 10.1016/J.CELREP.2022.110339.

Lee, C., Teng, Q., Zhong, R., Yuan, Y., Haghighat, M., and Ye, Z. H. (2012). Three Arabidopsis DUF579 domain-containing GXM proteins are methyltransferases catalyzing 4-O-methylation of glucuronic acid on xylan. Plant Cell Physiol. 53:1934–1949. 10.1093/PCP/PCS138.

Leitão, A. L., Costa, M. C., Gabriel, A. F., and Enguita, F. J. (2020). Interspecies communication in holobionts by non-coding RNA exchange. Int. J. Mol. Sci. 21:2333. 10.3390/ijms21072333.

Li, Y., Lu, Y. G., Shi, Y., Wu, L., Xu, Y. J., Huang, F., Guo, X. Y., Zhang, Y., Fan, J., Zhao, J. Q., et al. (2014). Multiple rice microRNAs are involved in immunity against the blast fungus Magnaporthe oryzae. Plant Physiol. 164:1077–1092. 10.1104/PP.113.230052.

Li, S. B., Xie, Z. Z., Hu, C. G., and Zhang, J. Z. (2016). A review of auxin response factors (ARFs) in plants. Front. Plant Sci. 7:175431. 10.3389/FPLS.2016.00047.

Li, L., Long, Y., Li, H., and Wu, X. (2020). Comparative transcriptome analysis reveals key pathways and hub genes in rapeseed during the early stage of Plasmodiophora brassicae infection. Front. Genet. 10:493966. 10.3389/FGENE.2019.01275.

Li, Q., Shah, N., Zhou, X., Wang, H., Yu, W., Luo, J., Liu, Y., Li, G., Liu, C., Zhang, C., et al. (2021). Identification of micro ribonucleic acids and their targets in response to Plasmodiophora brassicae infection in Brassica napus. Front. Plant Sci. 12:734419. 10.3389/fpls.2021.734419.

Liang, Y., Zhang, Y., Xu, L., Zhou, D., Jin, Z., Zhou, H., Lin, S., Cao, J., and Huang, L. (2019). CircRNA expression pattern and ceRNA and miRNA-mRNA networks involved in anther development in the CMS line of Brassica campestris. Int. J. Mol. Sci. 20:4808. 10.3390/ijms20194808.

Liao, Y., Smyth, G. K., and Shi, W. (2014). featureCounts: an efficient general purpose program for assigning sequence reads to genomic features. Bioinformatics 30:923–930. 10.1093/BIOINFORMATICS/BTT656.

Liao, R., Wei, X., Zhao, Y., Xie, Z., Nath, U. K., Yang, S., Su, H., Wang, Z., Li, L., Tian, B., et al. (2023). bra-miR167a targets ARF8 and negatively regulates Arabidopsis thaliana immunity against Plasmodiophora brassicae. Int. J. Mol. Sci. 24:11850. 10.3390/IJMS241411850/S1.

Llorente, F., Muskett, P., Sánchez-Vallet, A., López, G., Ramos, B., Sánchez-Rodríguez, C., Jordá, L., Parker, J., and Molina, A. (2008). Repression of the auxin response pathway increases Arabidopsis susceptibility to necrotrophic fungi. Mol. Plant 1:496–509. 10.1093/MP/SSN025.

Lombard, V., Golaconda Ramulu, H., Drula, E., Coutinho, P. M., and Henrissat, B. (2014). The carbohydrate-active enzymes database (CAZy) in 2013. Nucleic Acids Res. 42. 10.1093/NAR/GKT1178.

Love, M. I., Huber, W., and Anders, S. (2014). Moderated estimation of fold change and dispersion for RNA-seq data with DESeq2. Genome Biol. 15:550. 10.1186/s13059-014-0550-8.

Ludwig-Müller, J., and Schuller, A. (2008). What can we learn from clubroots: Alterations in host roots and hormone homeostasis caused by Plasmodiophora brassicae. Eur. J. Plant Pathol. 121:291–302. 10.1007/S10658-007-9237-2.

Luo, C., Bashir, N. H., Li, Z., Liu, C., Shi, Y., and Chu, H. (2024). Plant microRNAs regulate the defense response against pathogens. Front. Microbiol. 15:1434798. 10.3389/FMICB.2024.1434798/XML/NLM.

Makarewich, C. A., and Olson, E. N. (2017). Mining for Micropeptides 27. 10.1016/J.TCB.2017.04.006.

Martin, M. (2011). Cutadapt removes adapter sequences from high-throughput sequencing reads. EMBnet.journal 17.

Mehtab-Singh, Tripathi, R. K., Bekele, W. A., Tinker, N. A., and Singh, J. (2024). Differential expression and global analysis of miR156/SQUAMOSA promoter binding-like proteins (SPL) module in oat. Sci. Reports 2024 141 14:1–13. 10.1038/S41598-024-60739-7.

Middleton, H., Dozois, J. A., Monard, C., Daburon, V., Clostres, E., Tremblay, J., Combier, J.-P., Yergeau, É., and El Amrani, A. (2024). Rhizospheric miRNAs affect the plant microbiota 4. 10.1093/ISMECO/YCAE120.

Navarro, L., Dunoyer, P., Jay, F., Arnold, B., Dharmasiri, N., Estelle, M., Voinnet, O., and Jones, J. D. G. (2006). A plant miRNA contributes to antibacterial resistance by repressing auxin signaling 312:436–439.

Ormancey, M., Thuleau, P., Combier, J. P., and Plaza, S. (2023). The essentials on microRNA-encoded peptides from plants to animals. Biomol. 2023, Vol. 13, Page 206 13:206. 10.3390/BIOM13020206.

Ormancey, M., Guillotin, B., Ribeyre, C., Medina, C., Jariais, N., San Clemente, H., Thuleau, P., Plaza, S., Beck, M., and Combier, J. P. (2024). Immune-enhancing miPEPs reduce plant diseases and offer new solutions in agriculture. Plant Biotechnol. J. 22:13–15. 10.1111/PBI.14187.

Paul, P., Chhapekar, S. S., Rameneni, J. J., Oh, S. H., Dhandapani, V., Subburaj, S., Shin, S. Y., Ramchiary, N., Shin, C., Choi, S. R., et al. (2021). Mir1885 regulates disease tolerance genes in brassica rapa during early infection with plasmodiophora brassicae. Int. J. Mol. Sci. 22:9433. 10.3390/ijms22179433.

Pearson, W. R. (2016). Finding Protein and Nucleotide Similarities with FASTA. Curr. Protoc. Bioinforma. 53:3.9.1–3.9.25. 10.1002/0471250953.BI0309S53.

Penno, C., Clostres, E., Daburon, V., Chevallier, C., Durajova, Z., Dupont, A., Burel, A., Gauffre, F., Marchi, V., Combier, J.-P., et al. In prep. A pioneering investigation of secreted exosomes by root plants: RNA content and biological implications.

Petruschke, H., Schori, C., Canzler, S., Riesbeck, S., Poehlein, A., Daniel, R., Frei, D., Segessemann, T., Zimmerman, J., Marinos, G., et al. (2021). Discovery of novel community-relevant small proteins in a simplified human intestinal microbiome. Microbiome 9:1–19. 10.1186/S40168-020-00981-Z.

Prel, A., Dozier, C., Combier, J. P., Plaza, S., and Besson, A. (2021). Evidence that regulation of pri-miRNA/miRNA expression is not a general rule of miPEPs function in humans. Int. J. Mol. Sci. 2021, Vol. 22, Page 3432 22:3432. 10.3390/IJMS22073432.

R Core Team (2022). R: A language and environment for statistical computing. R Foundation for Statistical Computing Advance Access published 2022.

Rani, A., Singh, S., Yadav, P., Arora, H., Kaur, I., and Dhaka, N. (2023). MicroRNAs for understanding and improving agronomic traits in oilseed Brassicas. Plant Gene 34:100422. 10.1016/J.PLGENE.2023.100422.

Rhoades, M. W., Reinhart, B. J., Lim, L. P., Burge, C. B., Bartel, B., and Bartel, D. P. (2002). Prediction of plant microRNA targets. Cell 110:513–520. 10.1016/S0092-8674(02)00863-2.

Rice, P., Longden, L., and Bleasby, A. (2000). EMBOSS: The European Molecular Biology Open Software Suite. Trends Genet. 16:276–277. 10.1016/S0168-9525(00)02024-2.

Robin, A. H. K., Saha, G., Laila, R., Park, J. I., Kim, H. T., and Nou, I. S. (2020). Expression and role of biosynthetic, transporter, receptor, and responsive genes for auxin signaling during clubroot disease development. Int. J. Mol. Sci. 2020, Vol. 21, Page 5554 21:5554. 10.3390/IJMS21155554.

Rolfe, S. A., Strelkov, S. E., Links, M. G., Clarke, W. E., Robinson, S. J., Djavaheri, M., Malinowski, R., Haddadi, P., Kagale, S., Parkin, I. A. P., et al. (2016). The compact genome of the plant pathogen Plasmodiophora brassicae is adapted to intracellular interactions with host Brassica spp. BMC Genomics 17:1–15. 10.1186/S12864-016-2597-2.

Royston, J. P. (1982). An extension of Shapiro and Wilk’s W test for normality to large samples. Appl. Stat. 31:115. 10.2307/2347973.

Sang, Q., Fan, L., Liu, T., Qiu, Y., Du, J., Mo, B., Chen, M., and Chen, X. (2023). MicroRNA156 conditions auxin sensitivity to enable growth plasticity in response to environmental changes in Arabidopsis 14:1449. 10.1038/S41467-023-36774-9.

Schwab, R., Palatnik, J. F., Riester, M., Schommer, C., Schmid, M., and Weigel, D. (2005). Specific effects of microRNAs on the plant transcriptome. Dev. Cell 8:517–527. 10.1016/J.DEVCEL.2005.01.018.

Schwelm, A., Fogelqvist, J., Knaust, A., Jülke, S., Lilja, T., Bonilla-Rosso, G., Karlsson, M., Shevchenko, A., Dhandapani, V., Choi, S. R., et al. (2015). The Plasmodiophora brassicae genome reveals insights in its life cycle and ancestry of chitin synthases. Sci. Reports 2015 51 5:1–12. 10.1038/SREP11153.

Segonzac, C., and Monaghan, J. (2019). Modulation of plant innate immune signaling by small peptides. Curr. Opin. Plant Biol. 51:22–28. 10.1016/J.PBI.2019.03.007.

Song, S., Xu, Y., Huang, D., Ashraf, M. A., Li, J., Hu, W., Jin, Z., Zeng, C., Tang, F., Xu, B., et al. (2018). Identification and characterization of miRNA169 family members in banana (Musa acuminata L.) that respond to fusarium oxysporum f. sp. cubense infection in banana cultivars. PeerJ 6. 10.7717/PEERJ.6209.

Sun, T., Zhou, Q., Zhou, Z., Song, Y., Li, Y., Wang, H. Bin, and Liu, B. (2022). SQUINT positively regulates resistance to the pathogen Botrytis cinerea via miR156–SPL9 module in Arabidopsis. Plant Cell Physiol. 63:1414–1432. 10.1093/PCP/PCAC042.

Sweeney, B. A., Petrov, A. I., Ribas, C. E., Finn, R. D., Bateman, A., Szymanski, M., Karlowski, W. M., Seemann, S. E., Gorodkin, J., Cannone, J. J., et al. (2021). RNAcentral 2021: secondary structure integration, improved sequence search and new member databases. Nucleic Acids Res. 49:D212–D220. 10.1093/NAR/GKAA921.

Tarver, J. E., Donoghue, P. C. J., and Peterson, K. J. (2012). Do miRNAs have a deep evolutionary history? BioEssays 34:857–866. 10.1002/BIES.201200055.

Tu, J., Qin, L., Karunakaran, C., Wei, Y., and Peng, G. (2024). Lignin accumulation in cell wall plays a role in clubroot resistance. Front. Plant Sci. 15:1401265. 10.3389/FPLS.2024.1401265.

Uddin, M. N., Akhter, S., Chakraborty, R., Baek, J. H., Cha, J. Y., Park, S. J., Kang, H., Kim, W. Y., Lee, S. Y., Mackey, D., et al. (2017). SDE5, a putative RNA export protein, participates in plant innate immunity through a flagellin-dependent signaling pathway in Arabidopsis. Sci. Reports 2017 *71* 7:1–11. 10.1038/S41598-017-07918-X.

Vañó, M. S., Nourimand, M., MacLean, A., and Pérez-López, E. (2023). Getting to the root of a club - Understanding developmental manipulation by the clubroot pathogen. Semin. Cell Dev. Biol. 148–149:22–32. 10.1016/J.SEMCDB.2023.02.005.

Vazquez, F., Gasciolli, V., Crété, P., and Vaucheret, H. (2004). The nuclear dsRNA binding protein HYL1 is required for microRNA accumulation and plant development, but not posttranscriptional transgene silencing. Curr. Biol. 14:346–351. 10.1016/J.CUB.2004.01.035.

Verma, S. S., Rahman, M. H., Deyholos, M. K., Basu, U., and Kav, N. N. V. (2014). Differential expression of miRNAs in Brassica napus root following infection with Plasmodiophora brassicae. PLoS One 9:e86648. 10.1371/journal.pone.0086648.

Walerowski, P., Gündel, A., Yahaya, N., Truman, W., Sobczak, M., Olszak, M., Rolfe, S., Borisjuk, L., and Malinowski, R. (2018). Clubroot disease stimulates early steps of phloem differentiation and recruits SWEET sucrose transporters within developing galls. Plant Cell 30:3058–3073. 10.1105/TPC.18.00283.

Wang, H., and Wang, B. (2016). Extracellular vesicle microRNAs mediate skeletal muscle myogenesis and disease. Biomed. reports 5:296–300. 10.3892/br.2016.725.

Wang, D., Pajerowska-Mukhtar, K., Culler, A. H., and Dong, X. (2007). Salicylic acid inhibits pathogen growth in plants through repression of the auxin signaling pathway. Curr. Biol. 17:1784–1790. 10.1016/J.CUB.2007.09.025.

Wang, B., Sun, Y., Song, N., Zhao, M., Liu, R., Feng, H., Wang, X., and Kang, Z. (2017). Puccinia striiformis f. sp. tritici microRNA-like RNA 1 (Pst-milR1), an important pathogenicity factor of Pst, impairs wheat resistance to Pst by suppressing the wheat pathogenesis-related 2 gene. New Phytol. 215:338–350. 10.1111/nph.14577.

Wang, X., Cao, Y., Yang, J., Zhang, T., Yang, Q., Zhang, Y., Wang, D., and Cao, X. (2022). Transcription factor SmSPL2 inhibits the accumulation of salvianolic acid B and influences root architecture. Int. J. Mol. Sci. 23:13549. 10.3390/IJMS232113549/S1.

Wei, X., Liao, R., Zhang, X., Zhao, Y., Xie, Z., Yang, S., Su, H., Wang, Z., Zhang, L., Tian, B., et al. (2023). Integrative transcriptome, miRNAs, degradome, and phytohormone analysis of Brassica rapa L. in response to Plasmodiophora brassicae. Int. J. Mol. Sci. 24:2414. 10.3390/IJMS24032414/S1.

Weiberg, A., and Jin, H. (2015). Small RNAs — the secret agents in the plant–pathogen interactions. Curr. Opin. Plant Biol. 26:87–94. 10.1016/J.PBI.2015.05.033.

Weiberg, A., Wang, M., Bellinger, M., and Jin, H. (2014). Small RNAs: a new paradigm in plant-microbe interactions. Annu. Rev. Phytopathol. 52:495–516. 10.1146/ANNUREV-PHYTO-102313-045933.

Wickham, H. (2016). ggplot2: Elegant graphics for data analysis. Springer Verlag. 10.1007/978-3-319-24277-4.

Wu, G., Park, M. Y., Conway, S. R., Wang, J. W., Weigel, D., and Poethig, R. S. (2009). The sequential action of miR156 and miR172 regulates developmental timing in Arabidopsis. Cell 138:750–759. 10.1016/J.CELL.2009.06.031.

Xie, S., Jiang, H., Xu, Z., Xu, Q., and Cheng, B. (2017). Small RNA profiling reveals important roles for miRNAs in Arabidopsis response to Bacillus velezensis FZB42. Gene 629:9–15. 10.1016/J.GENE.2017.07.064.

Xu, M., Hu, T., Zhao, J., Park, M. Y., Earley, K. W., Wu, G., Yang, L., and Poethig, R. S. (2016). Developmental functions of miR156-regulated SQUAMOSA PROMOTER BINDING PROTEIN-LIKE (SPL) genes in Arabidopsis thaliana. PLOS Genet. 12:e1006263. 10.1371/JOURNAL.PGEN.1006263.

Yang, X., Zhang, L., Yang, Y., Schmid, M., and Wang, Y. (2021). miRNA mediated regulation and interaction between plants and pathogens. Int. J. Mol. Sci. 2021, Vol. 22, Page 2913 22:2913. 10.3390/IJMS22062913.

Yin, H., Hong, G., Li, L., Zhang, X., Kong, Y., Sun, Z., Li, J., Chen, J., and He, Y. (2019). MiR156/SPL9 regulates reactive oxygen species accumulation and immune response in arabidopsis thaliana. Phytopathology 109:632–642. 10.1094/PHYTO-08-18-0306-R.

Zhang, T., Zhao, Y. L., Zhao, J. H., Wang, S., Jin, Y., Chen, Z. Q., Fang, Y. Y., Hua, C. L., Ding, S. W., and Guo, H. S. (2016). Cotton plants export microRNAs to inhibit virulence gene expression in a fungal pathogen. Nat. Plants 2:1–6. 10.1038/nplants.2016.153.

Zhou, H., Lou, F., Bai, J., Sun, Y., Cai, W., Sun, L., Xu, Z., Liu, Z., Zhang, L., Yin, Q., et al. (2022). A peptide encoded by pri[miRNA[31 represses autoimmunity by promoting T reg differentiation. EMBO Rep. 23. 10.15252/EMBR.202153475.

